# Ubiquinone biosynthesis over the entire O_2_ range: characterization of a conserved, O_2_-independent pathway

**DOI:** 10.1101/587956

**Authors:** Ludovic Pelosi, Chau-Duy-Tam Vo, Sophie Saphia Abby, Laurent Loiseau, Bérengère Rascalou, Mahmoud Hajj Chehade, Bruno Faivre, Mathieu Goussé, Clothilde Chenal, Nadia Touati, Laurent Binet, David Cornu, Cameron David Fyfe, Marc Fontecave, Frédéric Barras, Murielle Lombard, Fabien Pierrel

**Author notes:** These authors contributed equally to the work (Ludovic Pelosi, Chau-Duy-Tam Vo).

## Abstract

Most bacteria can generate ATP by respiratory metabolism, in which electrons are shuttled from reduced substrates to terminal electron acceptors, via quinone molecules like *ubi*quinone. Dioxygen (O_2_) is the terminal electron acceptor of aerobic respiration and serves as a co-substrate in the biosynthesis of *ubi*quinone. Here, we characterize a novel, O_2_-independent pathway for the biosynthesis of *ubi*quinone. This pathway relies on three proteins UbiT (YhbT), UbiU (YhbU) and UbiV (YhbV). UbiT contains an SCP2 lipid-binding domain and is likely an accessory factor of the biosynthetic pathway, while UbiU-UbiV are involved in hydroxylation reactions and represent a novel class of O_2_-independent hydroxylases. We demonstrate that UbiU-UbiV form a heterodimer, wherein each protein binds a 4Fe-4S cluster via conserved cysteines that are essential for activity. The UbiT, -U, -V proteins are found in α-, β-, γ-proteobacterial clades including several human pathogens, supporting the widespread distribution of a previously-unrecognized capacity to synthesize *ubi*quinone in the absence of O_2_. Together, the O_2_-dependent and O_2_-independent *ubi*quinone biosynthesis pathways contribute to optimize bacterial metabolism over the entire O_2_ range.

**IMPORTANCE:** In order to colonize environments with large O_2_ gradients or fluctuating O_2_ levels, bacteria have developed metabolic responses that remain incompletely understood. Such adaptations have been recently linked to antibiotic resistance, virulence and the capacity to develop in complex ecosystems like the microbiota. Here, we identify a novel pathway for the biosynthesis of *ubi*quinone, a molecule with a key role in cellular bioenergetics. We link three uncharacterized genes of *Escherichia coli* to this pathway and show that the pathway functions independently from O_2_. In contrast, the long-described pathway for *ubi*quinone biosynthesis requires O_2_ as substrate. In fact, we find that many proteobacteria are equipped with the O_2_-dependent and O_2_-independent pathways, supporting that they are able to synthesize *ubi*quinone over the entire O_2_ range. Overall, we propose that the novel O_2_-independent pathway is part of the metabolic plasticity developed by proteobacteria to face varying environmental O_2_ levels.

## INTRODUCTION

Since the oxygenation of the Earth’s atmosphere some 2.3 billion years ago, many organisms adopted dioxygen (O_2_) as a terminal electron acceptor of their energy-producing respiratory chains [1]. Indeed, oxygenic (aerobic) respiration has a superior energetic output compared to anaerobic respiration or fermentation, which are both O_2_-independent processes [1]. In fact, several microorganisms, including many important human pathogens, are facultative anaerobes that are able to adopt either an aerobic or an anaerobic lifestyle depending on the environmental conditions [2,3]. In the laboratory, bacteria are usually cultured and studied under fully aerobic or completely anaerobic conditions (absence of O_2_) [4], whereas natural habitats cover the entire range of O_2_ concentrations [5]. For instance, large O_2_ gradients are typically encountered in the human large intestine, in biofilms or in transition zones between oxic and anoxic environments [5]. Moreover, bacteria can experience rapid transitions between environments with vastly different O_2_ contents, as during the infection process of enteric pathogens that progress along the gastrointestinal tract [3].

To maximize their bioenergetic capacities according to the various levels of O_2_ encountered in their environment, bacteria modulate the composition of their respiratory chains, notably the quinone species and the terminal reductases [3,4,6]. Quinones are lipophilic redox molecules that fuel electrons to terminal reductases, which reduce O_2_ whenever available, or alternative electron acceptors for instance nitrate, dimethyl sulfoxide (DMSO), trimethylamine N-oxide [7]. Naphtoquinones (menaquinone (MK) and demethylmenaquinone (DMK)) and *ubi*quinone (UQ) are the two main groups of bacterial quinones. (D)MK and UQ differ by the nature of their head group and the value of their redox midpoint potential [8]. (D)MK are considered “anaerobic quinones” since they function primarily in anaerobic respiration whereas UQ is considered an “aerobic quinone” because its supplies electrons mostly to the reductases that reduce O_2_ [1,8,9]. Accordingly, UQ is the main quinone of the facultative anaerobe *Escherichia coli* in aerobic conditions, whereas the naphtoquinones are predominant in the absence of O_2_ [10,11], UQ being nevertheless present.

The biosynthesis of UQ requires a total of eight reactions to modify the aromatic ring of the precursor, 4-hydroxybenzoic acid (4-HB): one prenylation, one decarboxylation, three hydroxylation and three methylation reactions (Figure 1A) [12]. In addition to the enzymes that catalyze the various steps, three accessory factors - UbiB, UbiJ and UbiK – are also needed. UbiB has an ATPase activity [13] and we showed that UbiJ and UbiK [14,15] belong to a multiprotein UQ biosynthesis complex, in which the SCP2 domain (sterol carrier protein 2) of UbiJ binds the hydrophobic UQ biosynthetic intermediates [16]. This UQ biosynthetic pathway is under the dependence of O_2_ since all three hydroxylases - UbiI, UbiH and UbiF – use O_2_ as a cosubstrate (Figure 1A) [17,18]. We showed recently that other hydroxylases, UbiL, UbiM and Coq7, replace UbiI, UbiH and UbiF in some proteobacteria [19]. The six hydroxylases have in common their dependence to O_2_ and thus function in UQ biosynthesis only when sufficient O_2_ is available. Interestingly, Alexander and Young established 40 years ago that *E. coli* was able to synthesize UQ anaerobically [20], suggesting the existence of an O_2_-independent biosynthesis pathway, which is still uncharacterized.

**Figure 1:**
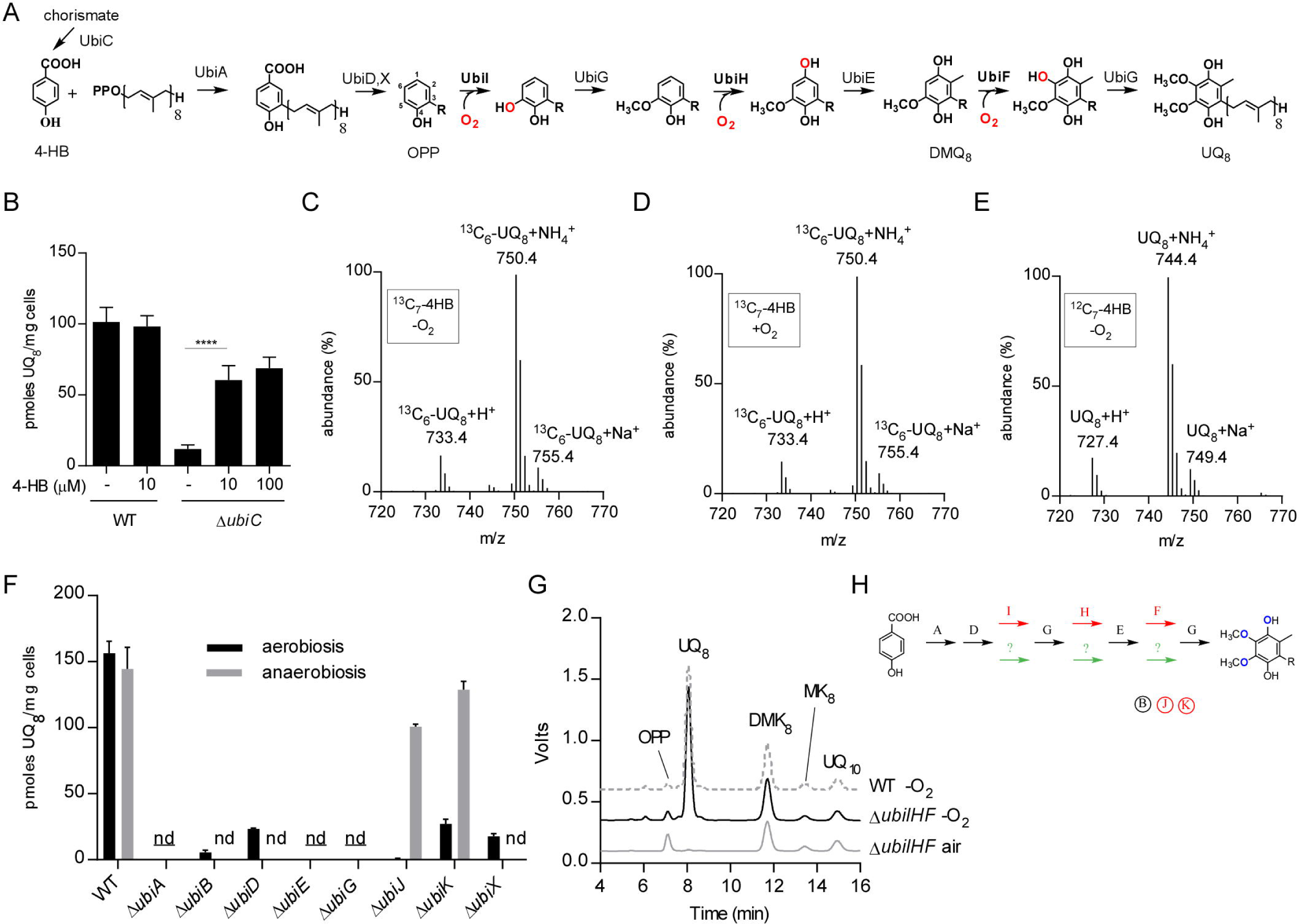
The aerobic and anaerobic UQ biosynthetic pathways differ only in the hydroxylation steps. A) O_2_-dependent UQ biosynthesis pathway in *E. coli*. The octaprenyl tail is represented by R on the biosynthetic intermediates and the numbering of the aromatic carbon atoms is shown on OPP. Abbreviations used are 4-HB for 4-hydroxybenzoic acid, OPP for octaprenylphenol, DMQ_8_ for C6-demethoxy-*ubi*quinone 8 and UQ_8_ for ubiquinone 8. B) UQ_8_ quantification of WT and Δ*ubi*C cells grown anaerobically in glycerol-nitrate medium supplemented or not with the indicated concentrations of 4-HB, mean ±SD (n=3-6), ****: p< 0.0001, unpaired Student’s t test. C-E) Mass spectra of UQ_8_ obtained by HPLC-MS analysis of lipid extracts from cells grown with ^13^C_7_-4HB either anaerobically (C) or aerobically (D), or anaerobically with unlabeled 4HB (E). F) UQ_8_ quantification from WT and Δ*ubi* cells grown anaerobically in SMGN medium overnight or aerobically in LB medium until OD 0.8. <u>nd</u> = not detected in aerobic and anaerobic conditions, nd= not detected in either aerobic or anaerobic conditions, mean ±SD (n=3-4). G) HPLC-ECD analyses (mobile phase 1) of lipid extracts from 1 mg of WT or Δ*ubi*IHF cells grown in LB medium in air or anaerobic conditions (-O_2_). Chromatograms are representative of n=3 independent experiments (UQ_10_ used as standard). H) UQ biosynthesis represented with Ubi enzymes specific to the O_2_-dependent pathway (red), to the O_2_-independent pathway (green), or common to both pathways (black). The same color code applies to the accessory factors (circled).

In this study, we describe the O_2_-independent UQ biosynthetic pathway in *E. coli* and identify three essential components, the UbiT, UbiU and UbiV proteins, formerly called YhbT, YhbU and YhbV. We show that the O_2_-independent UQ biosynthetic pathway is widely conserved in proteobacteria. UbiT likely functions as an accessory factor in the O_2_-independent UQ biosynthetic pathway and we show that UbiU and UbiV are involved in at least one O_2_-independent hydroxylation reaction. Moreover, we demonstrate that both UbiU and UbiV bind a [4Fe-4S] cluster essential for activity, which identifies these proteins as prototypes of a new class of O_2_-independent hydroxylases. Our results highlight that many proteobacterial species use two different and complementary molecular pathways to produce UQ over the entire continuum of environmental O_2_.

## RESULTS

### 4-HB is the precursor of UQ synthesized in anaerobic conditions

4-HB is the precursor of UQ synthesized in aerobic conditions [21]. Accordingly, an *E. coli* Δ*ubiC* mutant impaired in 4-HB biosynthesis [22], is deficient in UQ_8_ and is complemented by addition of 4-HB to the growth medium [23]. In order to evaluate whether 4-HB is also the precursor of the O_2_-independent UQ biosynthetic pathway, we grew a Δ*ubiC* strain anaerobically. The Δ*ubiC* strain showed a diminished level of UQ_8_, which was partially recovered by supplementation with 4-HB (Figure 1B). Furthermore, we grew the Δ*ubiC* strain in a medium supplemented with ^13^C_7_-4HB and we analyzed the labelling of biosynthesized UQ_8_ by HPLC-mass spectrometry (MS). In cells grown in aerobic or anaerobic conditions, the labelled form of UQ (^13^C_6_–UQ_8_, m/z= 750.4 for the NH_4_^+^ adduct) represented 98.3% and 97.3% (±0.2%) of the total UQ_8_ pool, respectively (Figure 1C, 1D). As expected, Δ*ubiC* cells grown anaerobically with unlabeled 4-HB didn’t show any ^13^C_6_-UQ_8_ (Figure 1E). Altogether, these results establish that 4-HB is the precursor of the O_2_-independent UQ biosynthetic pathway.

### Ubi enzymes, except hydroxylases, are common to the aerobic and anaerobic UQ biosynthesis pathways

The above result suggests that the UQ biosynthetic pathways decorate 4-HB with the same chemical groups irrespective of the presence of environmental O_2_. Thus, we evaluated whether the enzymes of the aerobic pathway are also involved in the O_2_-independent pathway by measuring the UQ_8_ content of knock-out (KO) strains grown in aerobic and anaerobic conditions. Deletion of *ubiA, ubiE* or *ubiG* abrogated UQ_8_ biosynthesis in both conditions whereas Δ*ubiB*, Δ*ubiD* or Δ*ubiX* strains synthesized a limited amount of UQ_8_ but only in aerobic conditions (Figure 1F). In contrast, *ubiJ* and *ubiK* had no effect on UQ biosynthesis under anaerobic conditions (Figure 1F).

In aerobic conditions, the hydroxylation reactions are catalyzed by the flavin monooxygenases (FMOs) UbiF, UbiH and UbiI that use dioxygen as a co-substrate [17,18,24] (Figure 1A). We previously reported that cells deleted for a single O_2_-dependent hydroxylase (Δ*ubiF*, Δ*ubiI* or Δ*ubiH*) were deficient in UQ, when cultured in the presence of air, but synthesized UQ in anaerobic conditions [17], consistent with the existence of an alternative hydroxylation system in the O_2_-independent pathway [20]. Indeed, we confirmed that all three FMOs are dispensable for the O_2_-independent UQ biosynthetic pathway since a triple mutant Δ*ubiF* Δ*ubiI* Δ*ubiH* was deficient for UQ when grown in air, but synthesized wild-type (WT) level of UQ_8_ in anaerobic conditions (Figure 1G). Together, our results demonstrate that the O_2_-dependent and O_2_-independent UQ biosynthetic pathways share the enzymes involved in the prenylation (UbiA), decarboxylation (UbiX and UbiD) and methylation reactions (UbiE and UbiG), but differ by their hydroxylases and the accessory factors UbiJ and UbiK (Figure 1H).

### Identification of three genes required for UQ biosynthesis in anaerobic conditions

To identify genes involved in the O_2_-independent UQ biosynthetic pathway, we cultivated anaerobically a collection of ∼ 200 *E. coli* strains that contained deletions covering multiple ORFs [25,26] and we analyzed their UQ_8_ content by HPLC-electrochemical detection (HPLC-ECD). We found a complete absence of UQ_8_ in strains ME4561, ME5034 and ME4746 that carry deletions encompassing, *ubiE-ubiJ-ubiB-ubiD, ubiG*, and *ubiX*, respectively (Table S1). Several other strains had a low UQ_8_ content and a poor growth in synthetic medium supplemented with glycerol and nitrate (SMGN). However, those strains showed better growth and higher UQ_8_ content in LB medium (Table S1). Thus we did not investigate them further, as a genetic defect affecting directly the O_2_-independent UQ pathway was unlikely. In contrast, ME4641 showed a profound UQ_8_ deficiency and robust anaerobic growth in LB and SMGN media (Table S1). Importantly, ME4641 had a WT UQ_8_ level when grown aerobically, suggesting that only the O_2_-independent pathway was altered (Figure 2A). ME4641 contains a deletion named OCL30-2 that covers 9 genes, 5 of them lacking an identified function (Figure 2B). To find the candidate gene involved in the anaerobic biosynthesis of UQ, we obtained 8 single gene KO strains from the Keio collection [27] and analyzed their quinone content after anaerobic growth (Figure 2C). The Δ*yhbT* and Δ*yhbU* strains were strongly deficient in UQ_8_. We then transduced the Δ*yhbT* and Δ*yhbU* mutations from the Keio strains into a MG1655 genetic background and also constructed the Δ*yhbV* strain, which was not available in the Keio collection. We found that all three strains had very low levels of UQ_8_ when grown in anaerobic conditions but showed normal levels after aerobic growth (Figure 2D, Table 1). In addition, the mutant strains showed a two-fold decrease in MK_8_ and a two-fold increase in DMK_8_ after anaerobic growth (Table 1). This effect might indirectly result from the UQ_8_ deficiency as it was also observed in the Δ*ubiG* strain (Figure S1A).

**Table 1:**
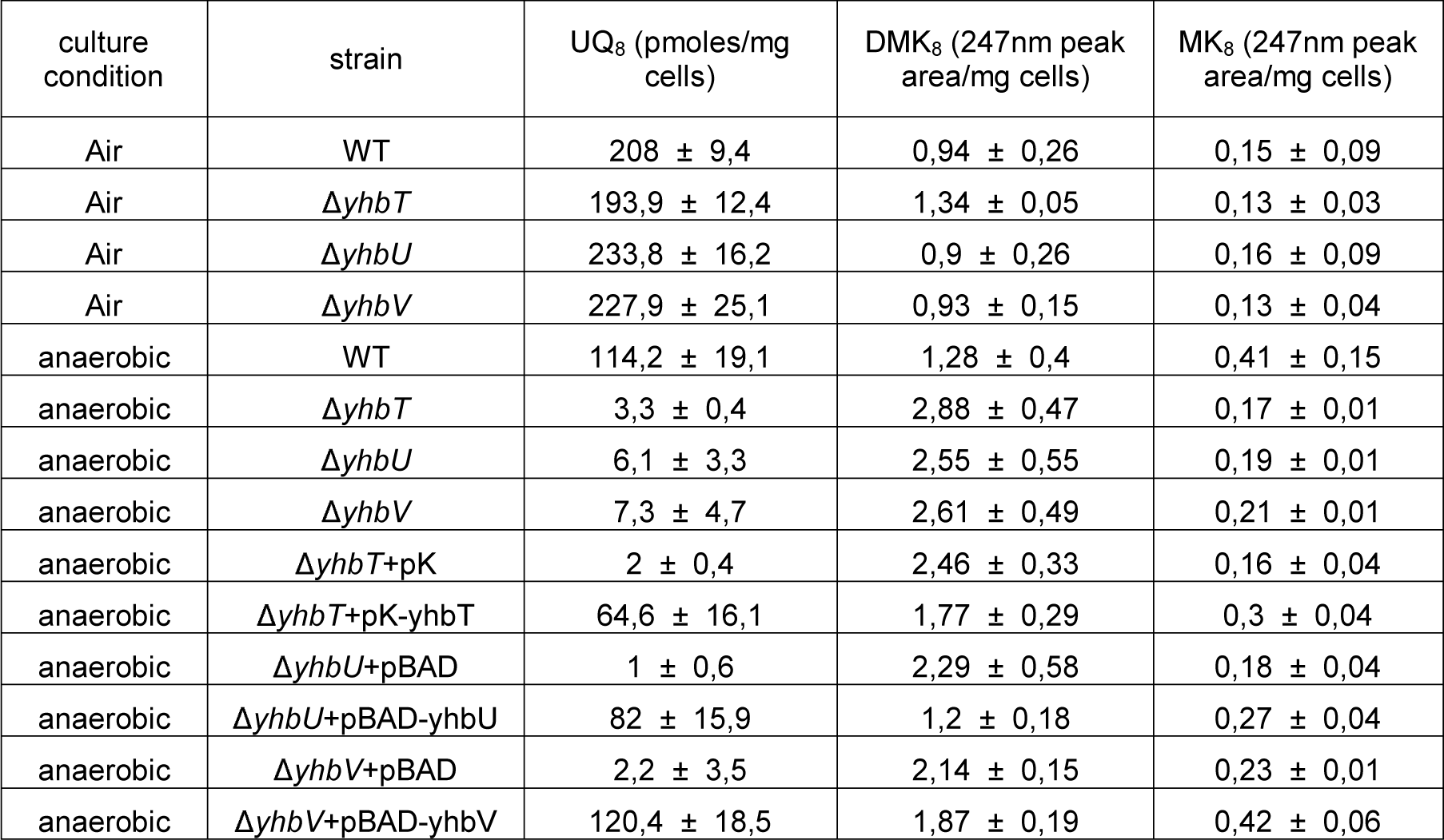
Quinone content of WT and Δ*yhb* strains cultured in SMGN either aerobically or anaerobically, n=3-5. For strains containing pK or pBAD vectors, SMGN was supplemented with 0.02% arabinose.

**Figure 2:**
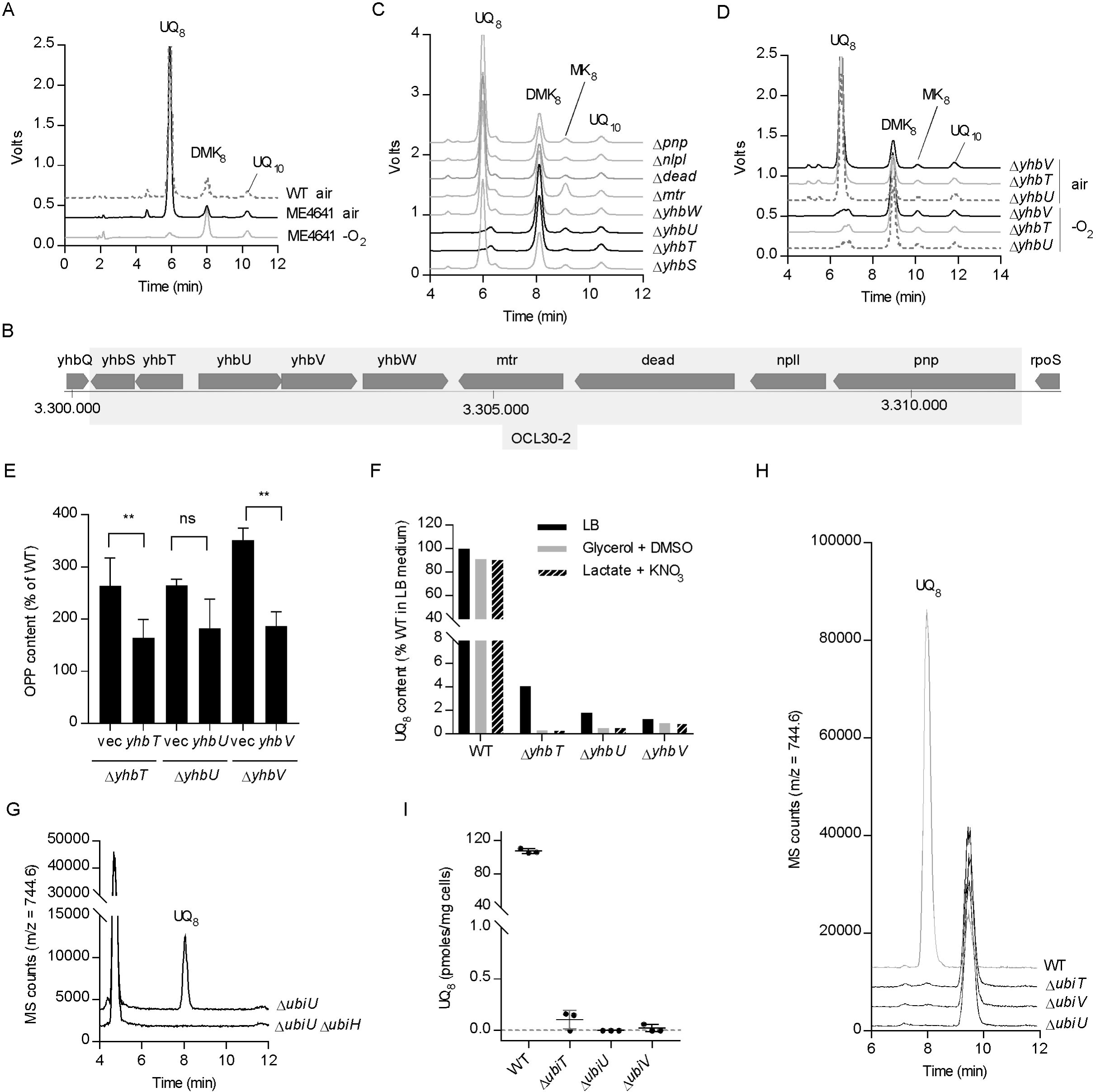
*yhbT, yhbU* and *yhbV* are essential to the anaerobic biosynthesis of UQ. A) HPLC-ECD analysis of lipid extracts from ME4641 strain grown in SMGN either aerobically or anaerobically (-O_2_). B) Genomic region covered by the OCL30-2 deletion in the ME4641 strain. C) HPLC-ECD analysis of lipid extracts from knock-out strains of the individual genes covered by the OCL30-2 deletion grown in SMGN anaerobically. D) HPLC-ECD analysis of lipid extracts from Δ*yhbT*, Δ*yhbU* and Δ*yhbV* strains constructed in the MG1655 background and grown in SMGN either aerobically or anaerobically. HPLC-ECD analyses with mobile phase 2 (A, C, D). E) OPP content (as % of WT, mass detection M+NH_4_^+^) in cells from table 2. The Δyhb strains contain either an empty plasmid or a plasmid carrying the indicated gene and were cultured anaerobically in SMGN containing 0.02% arabinose. Mean ±SD (n=3-5), **: p< 0.01, unpaired Student’s t test. F) UQ_8_ content (as % of WT grown in LB medium) of cells cultured anaerobically in SM containing the indicated carbon sources and electron acceptors. G) Single ion monitoring for UQ_8_(M+NH_4_^+^) in HPLC-MS analysis (mobile phase 1) of lipid extracts from 1 mg of Δ*ubiU* or Δ*ubiU* Δ*ubiH* cells grown in SMGN under anaerobic conditions. H) Single ion monitoring HPLC-MS analysis (mobile phase 1) of lipid extracts from 1.6 mg of cells grown in LB medium under strict anaerobic conditions and quenched in methanol. Chromatograms are representative of n=3 independent experiments (G,H). I) UQ_8_ content of cells described in H (quantification of the signal at 8 min with m/z= 744.6), mean ±SD (n=3).

### Deletion of *yhbT, yhbU* or *yhbV* causes UQ_8_ deficiency specifically in anaerobic conditions

We then transformed the mutant strains with an empty vector or a vector carrying a WT allele of the studied gene. In *yhb* KO strains expressing the corresponding gene from the plasmid, we observed a complementation of the UQ_8_ deficiency (Table 1) and a normalization of the levels of octaprenylphenol (OPP), an early UQ_8_ biosynthetic intermediate (Figure 2E). The ∼3 fold elevation of OPP in the *yhbT, -U, - V* KO mutants suggested that the O_2_-independent UQ biosynthetic pathway was blocked downstream of OPP in these strains. In these experiments, no cross-complementation was observed - for example the plasmid with *yhbV* had no effect in Δ*yhbT* or Δ*yhbU* strains - suggesting the absence of redundancy in the function of each gene. We also measured a profound UQ_8_ deficiency when the Δ*yhbT*, Δ*yhbU* and Δ*yhbV* strains were grown anaerobically in various media (glycerol + DMSO, lactate + KNO_3_) (Figure 2F), showing that the UQ_8_ biosynthetic defect is not linked to a particular carbon source or electron acceptor. Altogether, our results demonstrate that the *yhbT, yhbU* and *yhbV* genes are part of the O_2_-independent UQ biosynthetic pathway, so we propose to rename them *ubiT, ubiU* and *ubiV*, respectively.

**Table 2:** Characterization of UbiV proteins and UbiU-UbiV heterodimeric complexes (wild-type and mutants): metal content, and UV-Vis properties after attempts of anaerobic reconstitution of their Fe-S centers.

| proteins | $A_{280}/A_{410}$ | Iron content<br>(nmoles/nmole of protein) | Sulfur content<br>(nmoles/nmole of protein) |
| --- | --- | --- | --- |
| UbiV WT | 5,8 | $3.9 \pm 0.13$ | $3.3 \pm 0.18$ |
| UbiV C193A C197A | 8,4 | $2.4 \pm 0.02$ | $2.2 \pm 0.13$ |
| UbiV C39A C193A C197A | 15,5 | $1.9 \pm 0.01$ | $1.5 \pm 0.4$ |
| UbiV C180A C193A C197A | 26,7 | $0.7 \pm 0.05$ | $0.6 \pm 0.1$ |
| UbiV C39A C180A C193A C197A | 57,4 | $0.1 \pm 0.04$ | 0 |
| UbiU WT - UbiV WT | 4,3 | $7.9 \pm 0.05$ | $7.5 \pm 0.06$ |
| UbiU C169A C176A - UbiV WT | 7,3 | $4.7 \pm 0.03$ | $5.8 \pm 0.1$ |
| UbiU C193A C232A - UbiV WT | 8 | $3.9 \pm 0.05$ | $5.8 \pm 0.1$ |

### Complete absence of UQ biosynthesis in Δ*ubiT*, Δ*ubiU* or Δ*ubiV* mutants grown under strict anaerobic conditions

As the Δ*ubiT*, Δ*ubiU* or Δ*ubiV* strains still contained small amounts of UQ_8_ after growth in anaerobic conditions (Table 1, Figure 2F), we wondered how this UQ_8_ was synthesized. To verify if the O_2_-dependent pathway contributed to this synthesis, we inactivated the *ubiH* gene in the Δ*ubiU* strain. Sensitive HPLC-MS detection established that UQ_8_ was completely absent in extracts from the Δ*ubiH* Δ*ubiU* cells (Figure 2G). This result supported that the residual UQ_8_ synthesized in Δ*ubiU* cells originated from the O_2_-dependent pathway and suggested that our anaerobic media may contain traces amounts of O_2_ or that O_2_-dependent UQ biosynthesis may occur during the handling of cells, under normal atmosphere, prior to quinone extraction.

To eliminate the traces of O_2_, we took extra precautions in the degassing and inoculation of our media (see Material and Methods) and also added a reductant (L-Cysteine). In addition, we used the redox indicator resazurin to verify strict anaerobiosis during the entire culture. We also modified our sampling procedure to rapidly quench the anaerobic cells in ice-cold methanol, in order to prevent any O_2_-dependent UQ biosynthesis prior to quinone extraction. HPLC-MS analysis of extracts from cells cultivated and handled under such strict anaerobic conditions showed the nearly complete absence of UQ_8_ in Δ*ubiT*, Δ*ubiU* and Δ*ubiV* strains (Figure 2H, 2I). In contrast, the UQ_8_ level of the WT strain (Figure 2I) was comparable to those we measured previously in WT cells cultivated in suboptimal anaerobic conditions (Table 1) [14], establishing that UQ biosynthesis occurred independently from O_2_. Together, our results show that Δ*ubiT*, Δ*ubiU* and Δ*ubiV* cells are unable to synthesize UQ under strict anaerobic conditions unlike WT cells. Furthermore our results support that the low residual UQ content previously observed in Δ*ubiT*, Δ*ubiU* and Δ*ubiV* strains (Table 1, Figure 2F) resulted from the function of the O_2_-dependent pathway.

### *ubiT*,-*U*,-*V* strongly co-occur, exclusively in genomes with potential for UQ biosynthesis

We investigated the distribution of *ubiT*, -*U* and -*V* in a large genome dataset gathering 5750 genomes of bacteria and archaea. We found no evidence of genomes harboring matches for more than one of the three genes of interest outside of Alphaproteobacteria, Betaproteobacteria and Gammaproteobacteria (α-, β-, γ-proteobacteria) and three genomes of Acidithiobacillia (Table S2A). Interestingly, the three former classes are the only known so far to be able to produce UQ [1,8], consistent with a specific link between UbiT, -U, -V and UQ biosynthesis.

This was confirmed by analyzing in more details the distribution of *ubiT*, -*U* and -*V* and of three marker genes of the UQ biosynthetic pathway (*ubiA, ubiE* and *ubiG)* in 611 representative and reference genomes of α-, β-, γ-proteobacteria (Table S2B). We chose *ubiA, -G, -E* because they have a higher conservation compared to other genes like *ubiC, -D, -X* [28] and because they are part of the O_2_-dependent and O_2_-independent pathways (Figure 1H). 575 genomes had positive matches for *ubiA, -G, -E* and 589 for at least two of them. Regarding the distribution of *ubiT*, -*U* and -*V*, 221 genomes had matches for at least one of the three genes, and in 210 cases (95%), the three genes were present. Importantly, all genomes with *ubiT*, -*U* and -*V* harbored at least two of the three marker genes for the UQ biosynthetic pathway (Table S2B). In addition, we found that 22 out of the 29 proteobacterial orders analyzed had up to 50% genomes harboring a complete set of the *ubiT*, -*U* and -*V* genes (Figure 3A), demonstrating a wide taxonomic distribution. Overall, our analysis indicates a strong pattern of co-occurrence of *ubiT*, -*U* and -*V*, and demonstrates that they are uniquely found in genomes showing signs of UQ production.

**Figure 3:**
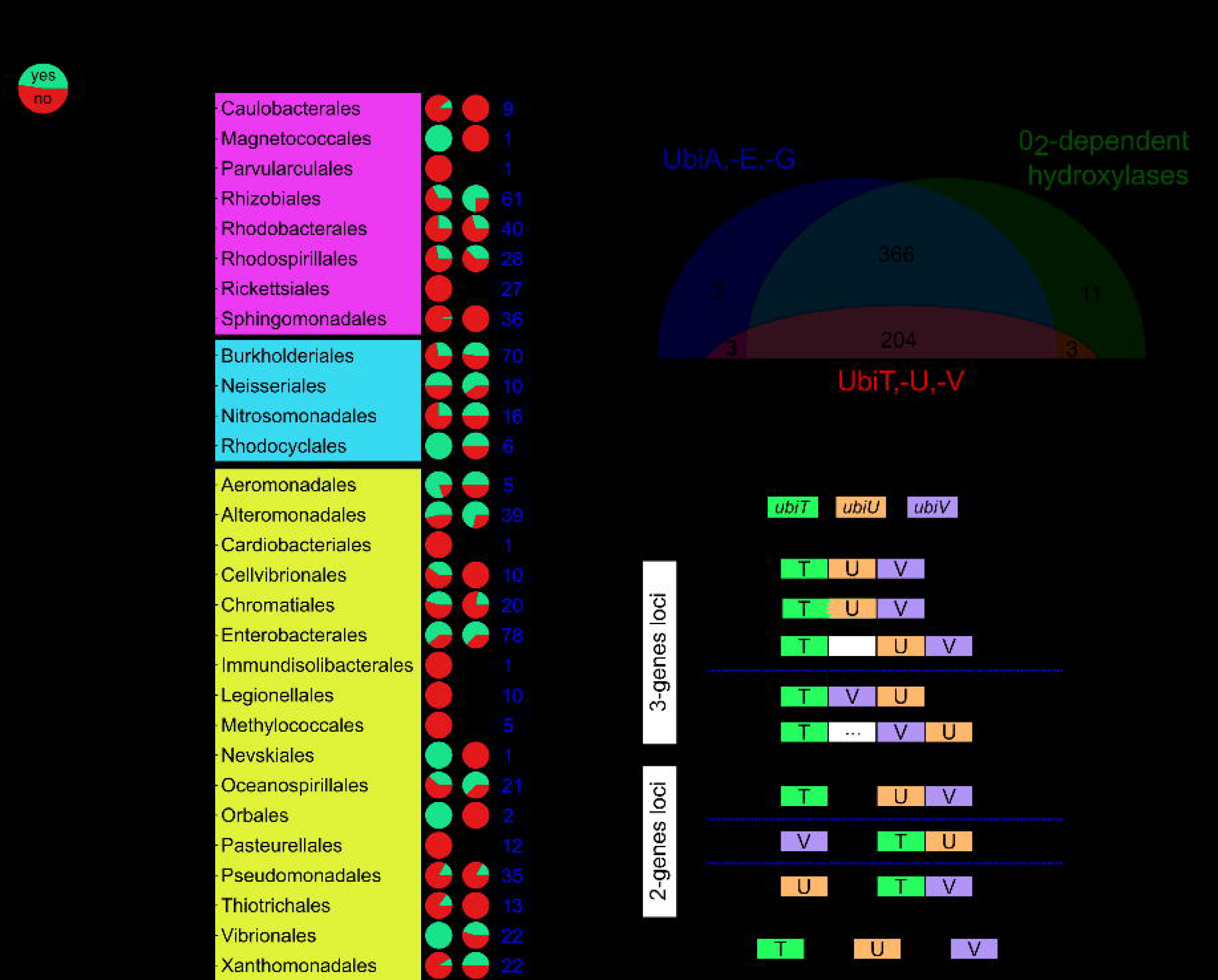
*ubiT, -U* and *-V* occurrence and genetic architecture in proteobacterial genomes. A) The proportion of genomes with (green) and without (red) all three genes “*ubiTUV*” (left column) is indicated for each proteobacterial order known to synthesize UQ. The middle column “3-genes loci” displays the proportion of genomes with the three genes either at a single locus (green) or at different loci (red). The number of genomes analyzed for each order is given in the right column (“Nb Genomes”). B) Occurrence in the reference proteobacterial genomes of the marker proteins (UbiA, -E, -G), of the O_2_– dependent hydroxylases and of the UbiT,-U, -V proteins. The number in bold represents the 3 genomes (*P. fulvum, M. marinus* and *O. formigenes*) containing exclusively the O_2_-independent pathway. C) The distinct genetic architectures found for *ubiT*, -U and -V in genomes where the three genes were present are displayed as boxes with different colors. Number of cases corresponding to each depicted architecture are given on the right. A white box corresponds to a gene found between the genes of interest, and a white box with dots corresponds to two to five genes between the genes of interest.

We found *ubiA, -G, -E* and O_2_-dependent hydroxylase genes in 570 genomes, of which 204 also contained *ubiT*, -*U* and –*V* (Table S2B and Figure 3B). Only 3 species, *Phaeospirillum fulvum, Magnetococcus marinus* and *Oxalobacter formigenes*, seem to rely exclusively on the O_2_-independent pathway for UQ production as their genomes contain the *ubiT*, -*U* and -*V* genes but no O_2_-dependent hydroxylases (Figure 3B). We noticed that the *ubiT*, -*U* and -*V* genes are found next to other *ubi* genes in *M. marinus* (Figure S1B). Interestingly, *P. fulvum* and *M. marinus* were described to synthesize UQ in microaerobic conditions [29-31] and *O. formigenes* has been documented as an obligate anaerobe [32]. Together, our results show that the O_2_-independent UQ biosynthesis pathway is widespread in α-, β-, γ-proteobacterial orders and co-exists with the O_2_-dependent pathway in 98% of the cases.

We then looked into the relative positioning of *ubiT*, -*U* and -*V* in the 210 genomes harboring the three genes. We found 106 cases, covering α-, β-, γ-proteobacterial orders, where they were encoded next to each other (“3-genes loci”) and 82 cases of a 2-genes locus with the third gene being encoded elsewhere in the genome (Figure 3C). Evaluation of the genetic architecture of *ubiT*, -*U* and –*V* revealed that *ubiU* and *ubiV* were found exactly next to each other in 69% of the loci (all 3-genes loci and in 39/82 of the 2-genes loci) and that the three genes were encoded in three separate parts of the genome only in 21 cases (Figure 3C). Interestingly, as an additional support for their involvement in a same function, we found an example of a gene fusion between *ubiT* and *ubiU* in two genomes from *Zymomonas mobilis* strains (α-proteobacteria), which also contain an *ubiV* gene directly downstream of the fused gene (Figure 3C).

### UbiT is an SCP2 protein and UbiU-V are required for the O2-independent hydroxylation of DMQ_8_

We then analyzed the sequences of the UbiT, UbiU and UbiV proteins. The major part of UbiT (amino acids 45-133, on a total of 174 aa in the *E. coli* protein) corresponds to a SCP2 domain (PFAM: PF02036, Figure S2). SCP2 domains typically form a hydrophobic cavity that binds various lipids [33] and our sequence alignment indeed showed the conservation of hydrophobic amino acids at several positions in the SCP2 domain of UbiT (Figure S2). Recently, we reported that UbiJ binds UQ biosynthetic intermediates in its SCP2 domain and organizes a multiprotein complex composed of several Ubi enzymes [16]. We propose that UbiT and its SCP2 domain may fulfill similar functions in the O_2_-independent UQ pathway, as UbiJ is required exclusively for the O_2_-dependent biosynthesis of UQ (Figure 1F).

UbiU and UbiV have ∼330 and 300 aa respectively, and contain an uncharacterized motif called peptidase U32 (PF01136) (Figure S3 and S4). Since only the hydroxylation reactions are uncharacterized in the O_2_-independent pathway (Figure 1H), we hypothesized that UbiU and UbiV might function in these steps. To test our hypothesis, we developed an *in vivo* assay based on the O_2_-independent conversion of labeled DMQ_8_ into labeled UQ_8_. This assay monitors the C6-hydroxylation and the subsequent O6-methylation (Figure 4A). Δ*ubiC* Δ*ubiF* cells grown aerobically with ^13^C_7_-4HB synthesized DMQ_8_, 73% of which was labelled with ^13^C_6_. Upon transfer to anaerobic conditions, the cells gradually converted a significant part of (^13^C_6_)-DMQ into (^13^C_6_)-UQ_8_ (Figure 4B). Inactivation of either *ubiU* or *ubiV* in Δ*ubiC* Δ*ubiF* cells did not perturb the accumulation of (^13^C_6_)-DMQ_8_ but prevented its conversion into (^13^C_6_)-UQ_8_ (Figure 4C-D). This result demonstrates that UbiU and UbiV are essential for the C6-hydroxylation reaction of the O_2_-independent UQ biosynthetic pathway.

**Figure 4:**
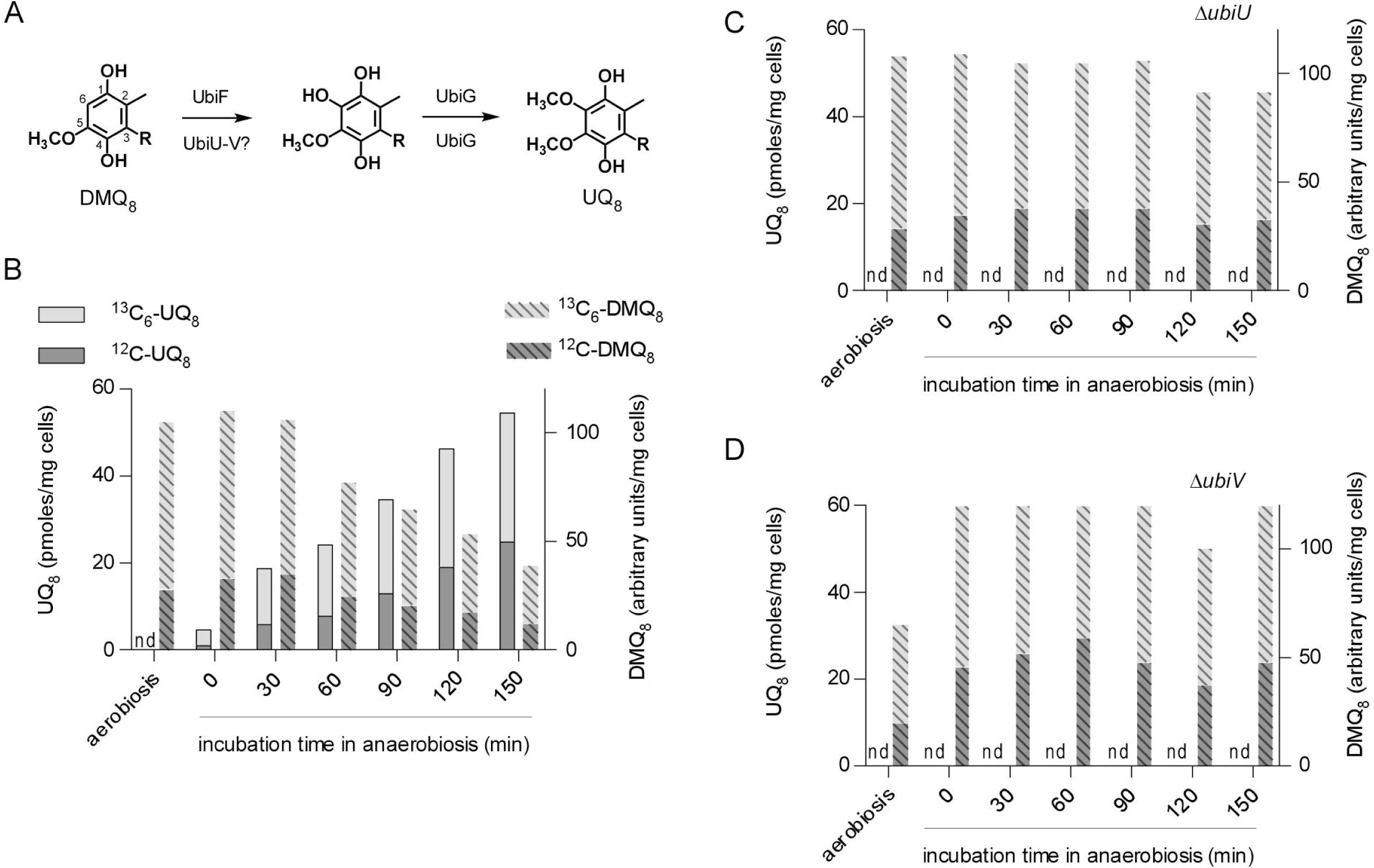
UbiU and UbiV are necessary for the anaerobic conversion of DMQ_8_ into UQ_8_. A) Conversion of DMQ_8_ to UQ_8_ with enzymes of the O_2_-dependent and the O_2_-independent pathways, indicated respectively above and below arrows (numbering of carbon atoms shown on DMQ_8_ and polyprenyl tail represented by R). B) Quantification by HPLC-MS (monitoring of Na^+^ adducts) of unlabeled (^12^C) and labeled (^13^C_6_) –DMQ_8_ and –UQ_9_ in Δ*ubi*C Δ*ubiF* cells after aerobic growth and transition to anaerobiosis. C-D) Same as in B with Δ*ubi*C Δ*ubiF* Δ*ubiU* cells (C) and Δ*ubiC* Δ*ubiF* Δ*ubiV* cells (D). nd, not detected, results representative of two independent experiments (B-D).

### UbiV contains a [4Fe-4S] cluster

To get insights into the potential presence of cofactors in UbiU and UbiV, we attempted to characterize them biochemically. UbiU was not soluble but we purified UbiV_6His_, which behaved as a monomer in solution (Figure S5A-B). UbiV was slightly pink-colored and had a UV-visible absorption spectrum with features in the 350-550 nm region (Figure 5A, dotted line), suggesting the presence of iron-sulfur (Fe-S) species [34-36]. We indeed detected substoichiometric amounts of iron and sulfur (0.2 Fe and 0.2 S/monomer), indicating a potential oxidative degradation of the Fe-S cluster during aerobic purification, as already observed with many other Fe-S proteins [37,38]. Consistent with this hypothesis, anaerobic reconstitution of the Fe-S cluster yielded a UbiV protein with 3.9 iron and 3.3 sulfur/monomer (Table 2) and with a UV-Vis spectrum characteristic of a [4Fe-4S]2+ cluster [39] (Figure 5A, solid line), that was affected by exposure to air (Figure S5C). The electron paramagnetic resonance (EPR) spectrum of the cluster reduced anaerobically displayed features characteristic of a [4Fe-4S]^1+^ cluster in the S = 1/2 state (Figure 5B) [38,40]. Overall, we conclude that, under anaerobic conditions, UbiV is able to bind one air-sensitive, redox-active [4Fe-4S] cluster.

**Figure 5:**
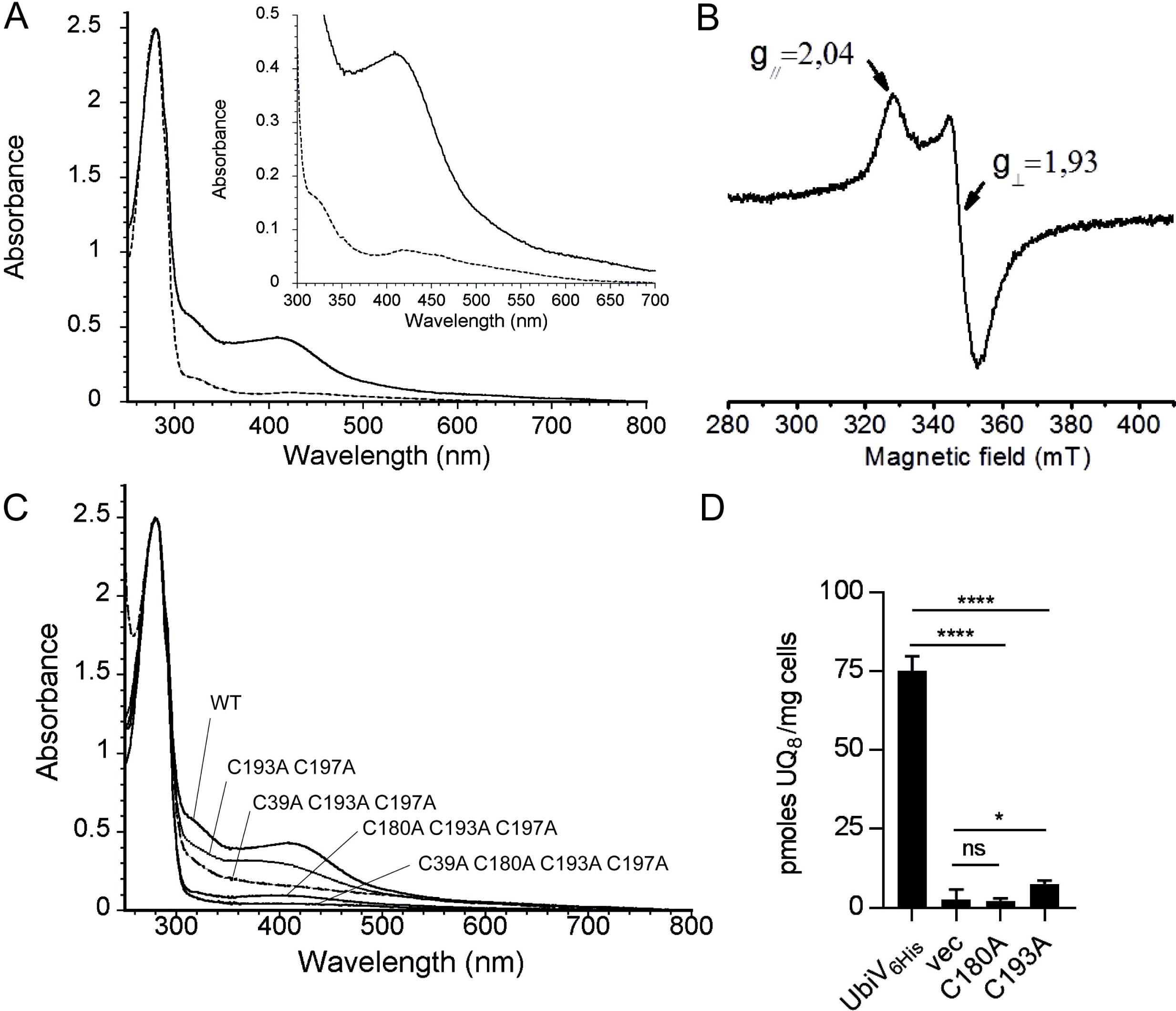
UbiV binds a [4Fe-4S] cluster. A) UV-visible absorption spectra of as-purified UbiV (dotted line, 47 µM) and reconstituted holo-UbiV (solid line, 41 µM); Inset: enlargement of the 300-700 nm region. The molar extinction coefficient ε_410nm_ was determined to be 10.8 ±_410nm_ 0.4 mM^-1^ cm^-1^ for holo-UbiV. B) X-band EPR spectrum of 785 µM dithionite-reduced holo-UbiV. Recording conditions: temperature, 10K; microwave power, 10 mW; modulation amplitude, 0.6 mT. C) Comparative UV-visible absorption spectra of WT and different Cys-to-Ala mutants of UbiV after Fe-S cluster reconstitution, with the following concentrations: 41 µM WT, 44 µM C193A C197A, 46 µM C39A C193A C197A, 47 µM C180A C193A C197A, and 54 µM C39A C180A C193A C197A. Proteins were in 50 mM Tris-HCl, pH 8.5, 25 mM NaCl, 15% glycerol, 1m M DTT (A-C). D) UQ_8_ quantification of Δ*ubiV* cells transformed with pBAD-UbiV_6His_, pBAD-UbiV_6His_ C180A, pBAD-UbiV_6His_ C193A or empty pBAD and grown overnight in anaerobic MSGN + 0.02% arabinose. Mean ±SD (n=4-5), *: p< 0.05, ****: p< 0.0001, unpaired Student’s t test.

Fe-S clusters are typically coordinated by cysteine residues [41,42] and we obtained evidence that the [4Fe-4S] cluster in UbiV is coordinated by four conserved cysteines arranged in a CX_n_CX_12_CX_3_C motif (Figure S4). Indeed, combinatorial elimination of C39, C180, C193, and C197 in double, triple and quadruple mutants resulted in proteins incapable of binding [Fe-S] clusters *in vivo*, as shown by the absence of absorption bands in the 350-550nm region of their UV-vis spectra (Figure S5D). Furthermore, after anaerobic reconstitution of the cluster, the Fe and S contents, and the absorbance at 410 nm were largely decreased in the double and triple mutants, and were undetectable in the quadruple mutant (Table 2 and Figure 5C). At last, we found that the mutation of C180 or C193 altered the function of UbiV *in vivo* (Figure 5D), suggesting that the [4Fe-4S] cluster is important for activity.

### UbiU contains a [4Fe-4S] cluster and forms a complex with UbiV

We succeeded purifying UbiU after coexpressing it with UbiV_6His_ (Figure S6A). The two proteins copurified in the form of a heterodimer (Figure S6A-B) that showed traces of Fe-S clusters (Figure 6A, dotted line), with substoichiometric amounts of iron and sulfide (0.4 Fe and 0.4 S/heterodimer). Reconstitution with iron and sulfide yielded a heterodimer with about 8 iron and 8 sulfur (Table 2) and a UV-visible spectrum characteristic of [4Fe-4S]2+ clusters (Figure 6A, solid line). The EPR spectrum of reduced UbiU-UbiV was also consistent with the presence of 2 different [4Fe-4S] clusters since it showed a composite signal, which reflects the presence of two different S=1/2 species (Figure 6B).

**Figure 6:**
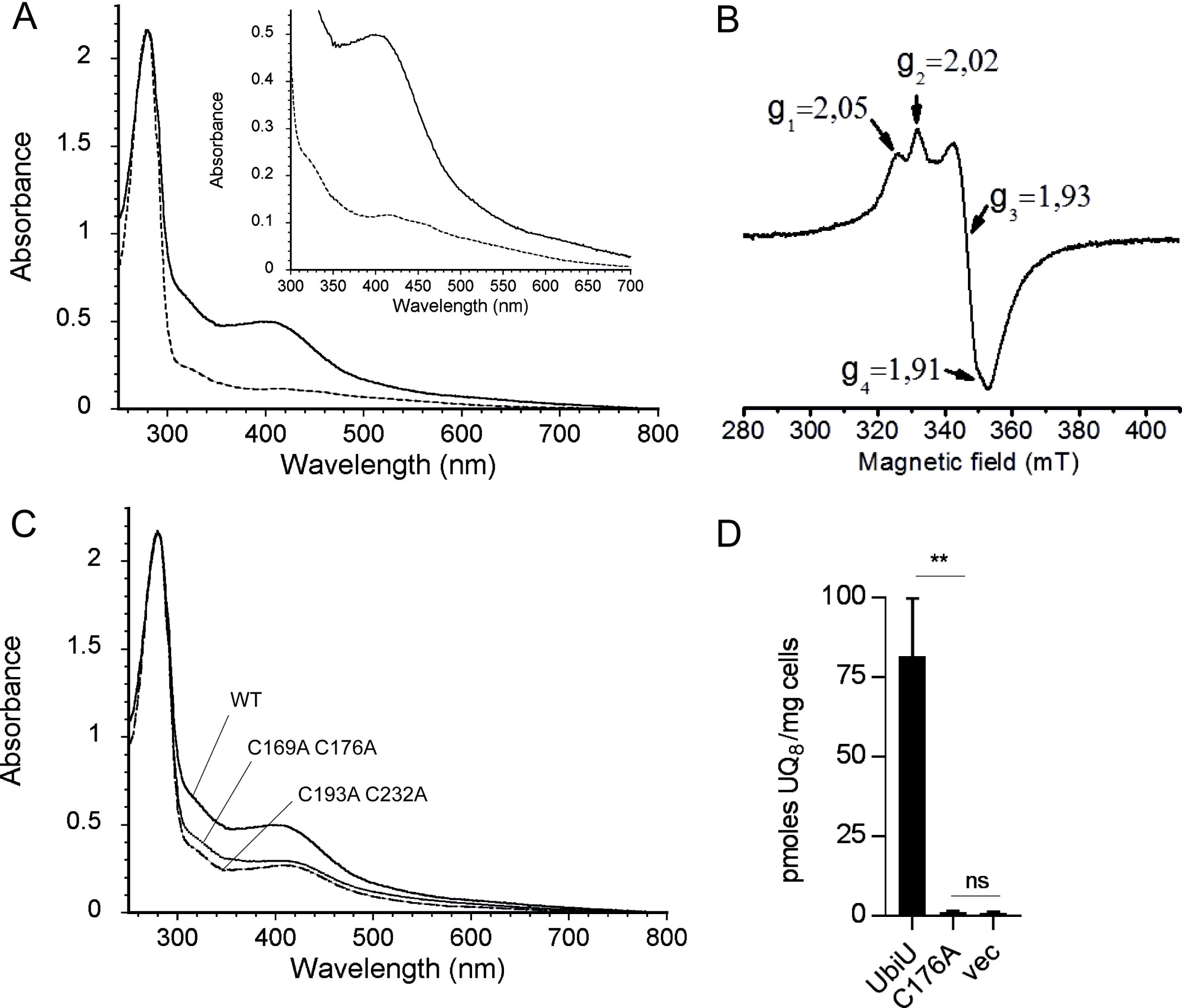
The UbiU-V complex binds two [4Fe-4S] clusters. A) UV-visible absorption spectra of as-purified UbiU-UbiV (dotted line, 17 µM) and reconstituted holo-UbiU-UbiV (solid line, 15.5 µM); Inset: enlargement of the 300-700 nm region. B) X-band EPR spectrum of 339 µM dithionite-reduced holo-UbiU-UbiV; Recording conditions: temperature, 10K; microwave power, 2 mW; modulation amplitude, 0.6 mT. C) Comparative UV-visible absorption spectra of Cys-to-Ala mutants of UbiU in the UbiU-UbiV complex after metal cluster reconstitution with the following concentrations: 15.5 µM WT, 16.0 µM UbiU-C169A C176A and 16.0 µM UbiU-C193A C232A. Proteins were in 50mM Tris-HCl, pH 8.5, 150mM NaCl, 15% glycerol and 1mM DTT (A-C). D) UQ_8_ quantification of Δ*ubiU* cells transformed with pBAD-UbiU (n=4), pBAD-UbiU C176A (n=2) or pBAD empty vector (n=3) and grown overnight in anaerobic MSGN+ 0.02% arabinose. Mean ±SD, **: p< 0.01, ns: not significant, unpaired Student’s t test.

Four strictly conserved cysteines are also found in UbiU (Figure S3) and we hypothesized that they might bind the [4Fe-4S]. We eliminated these cysteines in pairs and purified heterodimers composed of WT UbiV_6His_ and mutant UbiU (Figure S6C). After reconstitution, UbiU C169A C176A-UbiV and UbiU C193A C232A-UbiV had about half the iron as the WT heterodimer (Table 2), and their A_280_/A_410_ ratio were also diminished about two-fold (Figure 6C and Table 2). Altogether, our data clearly demonstrate that each protein of the heterodimeric UbiU-UbiV complex binds one [4Fe-4S] cluster and that the iron-chelating cysteines in UbiU are C169, C176, C193 and C232. Finally, an *in vivo* complementation assay demonstrated that C176 was important for the function of UbiU (Figure 6D).

### Many U32 proteases display motifs of four conserved cysteines

The presence of Fe-S clusters in UbiU and UbiV, two U32 proteases family members, led us to evaluate the presence of conserved Cys motifs in other U32 proteins. Kimura *et al.* reported a phylogenetic tree of 3521 peptidase U32 domains which formed 12 groups, belonging to 10 protein families [43]. We extracted and aligned the sequences of the 10 protein families and found highly conserved 4-cysteines clusters (97-100% conservation) in eight of them (Figure 7), suggesting an important functional role for these residues. Only families PepU32#5 and PepU32#6 had respectively none, and two mildly (60-80%) plus three poorly conserved cysteines in their sequences (40-65%). The cysteine motifs for each of the eight families showed a high degree of conservation, and strikingly, most of them could even be aligned with each other, the CX_6_CX_15_CX_3/4_C patterns appearing recurrently (Figure 7). To be noted, UbiV had a slightly distinct motif, with the first cysteine occurring much upstream of the three others and outside of the U32 domain. Overall, our data suggest that most members of the U32 peptidase family may contain a 4Fe-4S cluster coordinated by conserved cysteines, similar to what we demonstrated for UbiU and UbiV.

**Figure 7:**
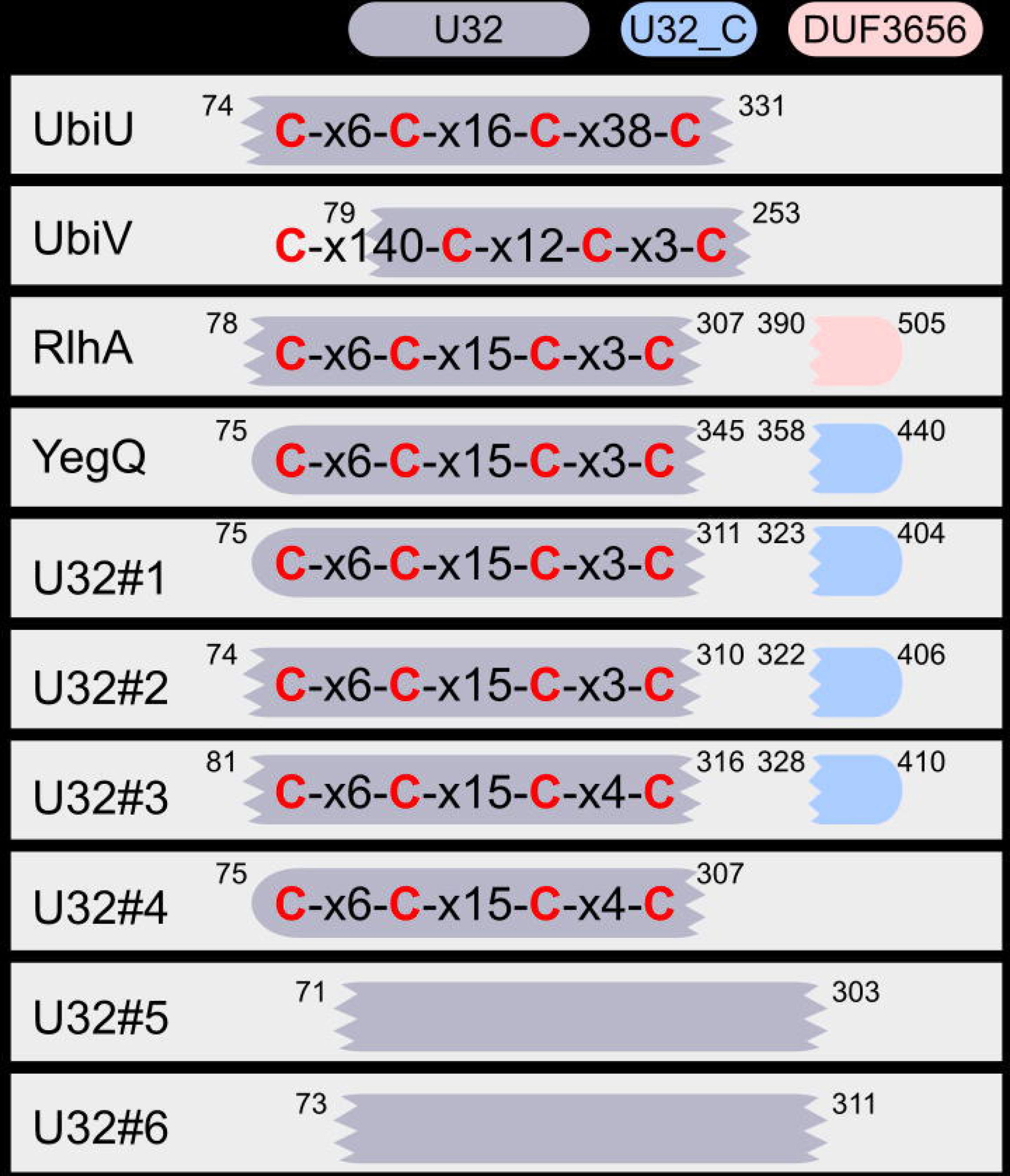
Conserved four cysteines motifs in the U32 protease family. The conserved 4-cysteines motifs and PFAM domains (colored boxes) found in each U32 protease family are displayed for a set of reference sequences. These motifs were obtained by aligning the sequences listed by Kimura *et al*. [43]. Conserved cysteines are in red, “x6” indicates that 6 residues were found in between two conserved cysteines. Positions of the domains are displayed on the outside of the boxes for the reference sequences. Scrambled extremities show interrupted match for the PFAM domain. No conserved cysteines were found for U32#5 and U32#6 (see main text). Reference sequences were from *E. coli* for UbiU, UbiV, YegQ and RhlA (YHBU_ECOLI, YHBV_ECOLI, YEGQ_ECOLI, YDCP_ECOLI for RlhA). For the rest of the families, the sequence accession numbers were: R7JPV1_9FIRM for U32#1, R6XKQ3_9CLOT for U32#2, S1NZZ5_9ENTE for U32#3, H1YXA1_9EURY for U32#4, H3NJ45_9LACT for U32#5, D5MIQ1_9BACT for U32#6.

## DISCUSSION

### The O_2_-independent UQ biosynthesis pathway is widespread in proteobacteria

The evidence for an O_2_-independent synthesis of UQ by *E. coli* was reported more than forty years ago [20], yet this pathway remained uncharacterized until now. Circumstantial evidence had been obtained that a few species – limited, to our knowledge, to *E. coli* [20], *Rhodobacter sphaeroides* [44], *Paracoccus denitrificans* [45], *Halorhodospira halophila* [46] – were able to synthesize UQ in anaerobic conditions, as demonstrated by biochemical measurements of the quinone content of cells grown anaerobically. Here, we demonstrate that the O_2_-dependent and O_2_-independent UQ biosynthesis pathways differ only by three hydroxylation steps (Figure 1), and we identify three genes, *ubiT,-U,-V* that are essential for the O_2_-independent biosynthesis of UQ in *E. coli* (Figure 2). The facts that the UbiT, -U, -V proteins are widespread in α, β, γ–proteobacterial clades (Figure 3) and co-occur with UbiA, -E, -G enzymes (Table S2B), reveal UbiT, -U, -V as key elements of the broadly-distributed, O_2_-independent UQ pathway. Overall, our data support that many proteobacteria have the previously-unrecognized capacity to synthesize UQ independently from O_2_.

### Physiological possibilities offered by UQ biosynthesis over the entire O_2_ range

In our set of reference genomes, only three species (*P. fulvum, M. marinus* and *O. formigenes*) seem to rely exclusively on the O_2_-independent pathway for UQ production (Table S2B). Indeed, the vast majority of proteobacteria with the O_2_-independent UQ biosynthesis pathway also possess the O_2_-dependent hydroxylases of the aerobic pathway (207 out of 210) (Figure 3B). This result supports that both pathways confer physiological advantages, allowing production of UQ over the entire spectrum of O_2_ levels encountered by facultative aerobes.

So-called microaerobes, able to respire O_2_ in microaerobic conditions, are very abundant in Nature [5] and *E. coli* is known to respire nanomolar O_2_ concentrations [47]. To sustain O_2_ respiration in the microaerobic range, these organisms are equipped with high affinity O_2_ reductases [5,47,48]. These enzymes reduce efficiently the low levels of environmental O_2_ present at the cell’s membrane, leaving the cytoplasm devoid of any O_2_ [49]. Under such conditions, UQ - which is the main electron donor for the high affinity O_2_ reductases *bd*I and *bd*II of *E. coli* [9] - must therefore be synthesized via the O_2_-independent pathway. Overall, we believe that the O_2_-independent UQ biosynthesis pathway operates not only in anaerobic conditions but also in microaerobic conditions, in which UQ is likely crucial for bacterial physiology.

The O_2_-independent UQ biosynthesis pathway may also confer a significant advantage to facultative bacteria in case of a rapid transition from an anaerobic to an aerobic environment. Indeed, anaerobic biosynthesis of UQ will result in cellular membranes containing UQ at the time of the transition, allowing an immediate switch to the energetically favorable metabolism of O_2_ respiration. Our identification of the anaerobic UQ pathway provides the unique opportunity to selectively disrupt UQ biosynthesis depending on O_2_ levels and should foster new research on bacterial physiology in the microaerobic range. Indeed, apart from *E. coli*, which was thoroughly studied over the microaerobic range [4,49,50], details on bacterial physiology in microaerobiosis are scarce.

### *ubiT,U,V* mutants and pathogenicity

In addition to bioenergetics *per se*, anaerobic and microaerobic respirations are thought to be important for pathogenicity [3,51]. Interestingly, homologs of UbiT, -U, -V have been linked to pathogenicity in several bacterial models. Indeed, the inactivation of *ubiU-V* homologs in *Proteus mirabilis* leads to a decreased infection of the urinary tract of mice [52] and to a diminished virulence of *Yersinia ruckeri* [53], a pathogen that develops in the gut of fish, an environment with a notoriously low O_2_ content. Furthermore, inactivation of PA3911 [54] (*ubiT*) and PA3912-PA3913 [55] (*ubiU-V*) in *Pseudomonas aeruginosa*, abolished nitrate respiration, the main anaerobic metabolism used by the bacterium in the lungs of cystic fibrosis patients [56,57]. Based on our results showing that the deletion of *ubiT, ubiU* or *ubiV* abrogates the O_2_-independent biosynthesis of UQ in *E. coli*, we suggest that the attenuation of the mutants discussed above results from their UQ deficiency in microaerobic / anaerobic conditions.

### Proposed roles for UbiT, UbiU and UbiV

UbiT possesses an SCP2 domain, similar to UbiJ that we recently demonstrated to be an accessory factor that binds the hydrophobic UQ biosynthetic intermediates and structures a multiprotein Ubi complex [16]. Since UbiJ functions exclusively in the O_2_-dependent pathway, whereas UbiT is important only for the O_2_-independent pathway, we propose that UbiT may fulfill, in anaerobiosis, the same functions than UbiJ in aerobiosis. Whether UbiT is part of a complex and is able to bind UQ biosynthetic intermediates, will be addressed in future studies. Interestingly, PA3911, the homolog of UbiT in *P. aeruginosa*, was recently shown to bind phosphatidic acid [54], demonstrating an affinity of UbiT for lipid molecules.

UbiU and UbiV form a tight heterodimer suggesting that the proteins function together, as further supported by the fact that the *ubiU* and *ubiV* genes co-occur in genomes in 99% of the cases and that they are mostly found next to each other (Figure 3). We demonstrated that UbiU and UbiV are both required for the O_2_-independent C6-hydroxylation of DMQ and the accumulation of OPP in Δ*ubiU* or Δ*ubiV* mutants suggests that the two proteins may also function in C5-hydroxylation. We want to emphasize our recent demonstration that a single hydroxylase catalyzes all three hydroxylation steps in the O_2_-dependent UQ pathway of *Neisseria meningitidis* [19]. This result showed that three different enzymes are not necessarily required and opens the possibility that UbiU-V may in fact catalyze all three hydroxylation reactions of the O_2_-independent UQ biosynthesis pathway. Establishing the hydroxylase activity and the regioselectivity of UbiU-V will require the development of an *in vitro* assay, a challenging task given that the oxygen donor of the reaction is currently unknown and that the substrates are not commercial and highly hydrophobic. Of note, one of the O_2_-dependent hydroxylases was shown to hydroxylate DMQ_0_, a substrate analog with no polyprenyl side chain [58], suggesting that an *in vitro* assay for UbiU-V might be developed with soluble analogs.

### Fe-S clusters in UbiU-V and other members of the U32 peptidase family

Up to now, members of the peptidase family U32 were not shown to bind Fe-S clusters or to contain any of the ~30 cysteine motifs found in well-characterized iron-sulfur proteins [59]. The expression, purification and spectroscopic characterization of UbiV and of the UbiU-UbiV heterodimeric complex clearly showed that each protein contains one [4Fe-4S] cluster (Figure 5 and Figure 6). Mutation of the candidate cysteine ligands, arranged in a CX_6_CX_16_CX_38_C motif in UbiU and in a CX_n_CX_12_CX_3_C motif in UbiV, disrupted Fe-S binding and abolished *in vivo* complementation, suggesting a crucial function of the [4Fe-4S] clusters in these proteins. The conservation of a CX_6_CX_15_CX_3/4_C motif in other U32 proteases supports that these proteins likely bind Fe-S clusters. This hypothesis should guide and stimulate investigations of U32 members, very few of which have currently an established molecular function ([60], http://www.ebi.ac.uk/merops/). Interestingly, RlhA, a member of the U32 proteases family involved in the C-hydroxylation of a cytidine on *E. coli* 23S rRNA, was recently shown to be connected to iron metabolism [43], corroborating our suggestion that RlhA is also an Fe-S protein.

In biological systems, Fe-S clusters are mainly known to be involved in electron transfer reactions, but also in substrate binding and activation, in transcription regulation, in iron storage, and as a sulfur donor [41,61-63]. The role of the Fe-S clusters in UbiU and UbiV is unknown at this stage. Our current working hypothesis is a role as an electron transfer chain between the substrate (the UQ biosynthetic intermediate to be hydroxylated) and an unidentified electron acceptor required for the activation of the substrate. Clearly, the Fe-S clusters of UbiU-V are distinct from the molybdenum cofactor present in molybdenum-containing hydroxylases, the only family currently known to catalyze O_2_-independent hydroxylation reactions [64]. Together, our results identify UbiU and UbiV as prototypes of a novel class of O_2_-independent hydroxylases and extend the framework of the chemically-fascinating O_2_-independent hydroxylation reactions.

## MATERIALS and METHODS

### Strain construction

Strains used in this study are listed in table S3. We obtained the collection of *E. coli* strains containing large and medium deletions from the National BioResource Project, National Institute of Genetics, Japan (http://www.shigen.nig.ac.jp/ecoli/pec/).

The Δ*ubiA*∷*cat*, Δ*ubiD*∷*cat*, Δ*ubiT*∷*cat* and Δ*ubi*V∷cat mutations were constructed in a one-step inactivation of *ubi* genes as described [65]. A DNA fragment containing the cat gene flanked with a 5⍰ and 3⍰ region bordering the *E.coli ubi* genes was amplified by PCR using pKD3 as a template and oligonucleotides 5wanner and 3wanner (Table S3). Strain BW25113, carrying the pKD46 plasmid, was transformed by electroporation with the amplified fragment and cat^R^ colonies were selected. The replacement of chromosomal *ubi* by cat gene was verified by PCR amplification in the cat^R^ clones. Mutations (including *ubiU∷kan* from the keio strain) were introduced into MG1655 strains by P1 *vir* transduction [66], selecting for the appropriate antibiotic resistance. The antibiotic resistance cassettes were eliminated when needed using plasmid pCP20 as described [67].

### Plasmid construction

All plasmids generated in this study were verified by DNA sequencing. The *yhbU, yhbT and yhbV* inserts (UniProtKB: P45527, P64599, P45475) were obtained by PCR amplification using *E. coli* MG1655 as template and the oligonucleotides pairs *yhbU5/yhbU3, yhbT5/yhbT3* and *yhbV5/yhbV3*, respectively (Table S3). *Yhb* inserts were EcoRI-SalI digested and inserted into EcoRI-SalI-digested pBAD24 plasmids, yielding the *pBAD-yhbU, pK-yhbT* or *pBAD-yhbV* plasmids, respectively.

To create a plasmid expressing the *ubi*V (*yhbV*) ORF as C-terminally His-tagged protein, the *ubiV* gene was amplified using pET-22-UbiV-FW (introducing the NdeI site) and pET-22-UbiV-RV (introducing the XhoI site) as primers and pBAD-*yhbV* as template. The NdeI and XhoI digested amplicon was ligated to NdeI and XhoI digested pET-22b(+) plasmid to obtain pET-22-UbiV.

The plasmid pETDUET-UbiUV containing UbiU in MCS1 and UbiV in MCS2 was obtained as follows. *ubiU* was amplified from pBAD-*yhbU* using pETDUET-UbiU-FW (introducing the NcoI site) and pETDUET-UbiU-RV (introducing the EcoRI site) as primers. The NcoI and EcoRI digested amplicon was ligated to NcoI and EcoRI digested pETDUET-1-plasmid to obtain pETDUET-UbiU. The *ubiV* gene was then cloned from pET-22-UbiV into the MSC2 of pETDUET-UbiU by PCR amplification with pET-22-UbiV-FW and pETDUET-UbiV-RV (introducing the C-terminal His_6_-tag) as primers. The NdeI and XhoI digested amplicon was ligated to NdeI and XhoI digested pETDUET-UbiU to obtain pETDUET-UbiUV.

A hexahistidine-tag was fused at the N-terminal extremity of UbiV to create pBAD-UbiV_6His_. The *ubiV6His* gene was obtained by PCR amplification (Phusion High-Fidelity DNA Polymerase) using pBAD-UbiV as a template and 6HisV5 (introducing the NcoI site) and 6HisV3 (introducing the HindIII site and the DNA sequence of 6His-tag) as primers. The NcoI/HindIII digested amplicon was cloned into the NcoI/HindII-digested pBAD plasmid.

Variants of UbiV and UbiU were constructed using the Q5 Site-Directed Mutagenesis Kit (New England Biolabs) according to the manufacturer’s specifications. The plasmids (pET-22b-*UbiV*, pETDUET-UbiUV, pBAD-*yhbU* or pBAD-*yhbV*) were used as templates in conjunction with the appropriate primers for each respective amino acid substitution.

### Culture conditions

*E. coli* strains were grown at 37 °C in lysogeny broth (LB) medium or in synthetic medium (SM) containing either 0.4% glycerol (w/v), 0.4% lactate (w/v) or 0.2% glucose (w/v) as carbon sources. Autoclaved SM medium was supplemented with 0.5% casaminoacids (w/v) and with 1/100 volume of a filter-sterile solution of 1mM CaCl_2_, 200 mM MgCl_2_, 1% thiamine (w/v) [68]. Ampicillin (50 mg/L), kanamycin (25 mg/L), and chloramphenicol (25 mg/L) were added from stocks (1000X solution sterilized through 0.22 µm filters and stored at −20°C), when needed. When needed, 0.02% arabinose was added to induce the expression of genes carried on pBAD and pK plasmids. External electron acceptors like KNO_3_ (100 mM) or dimethylsulfoxide (DMSO, 50 mM) were added to SM medium for anaerobic cultures. Anaerobic cultures were performed in Hungate tubes containing 12 mL medium deoxygenated by argon (O_2_<0.1 ppm) bubbling for 25 min before autoclave (in case of LB medium, 0.05% antifoam (Sigma) was added). Hungate tubes were inoculated through the septum with 100 µL of overnight precultures taken with disposable syringe and needles from closed Eppendorf tubes filled to the top. Aerobic cultures were performed in Erlenmeyer flasks filled to 1/10 of the maximal volume and shaken at 180 rpm.

For the initial screen, we grew ME strains anaerobically in SMGN (SM medium supplemented with glycerol and nitrate). Strains that presented a severe growth defect or a low UQ_8_ content were subsequently grown anaerobically in LB medium.

Cultures were cooled down on ice before transferring 5-10 mL volumes into 15 mL Falcon tubes for centrifugation at 3200 g, 4°C, 10 min. Cell pellets were washed in 1 mL ice-cold PBS and transferred to pre-weighted 1.5 mL Eppendorf tubes. After centrifugation at 12000 g, 4°C, 1 min and elimination of supernatant, the cells wet weight was determined (∼10-20 mg) and pellets were stored at −20°C prior to quinone extraction. We note that these steps were conducted under normal atmosphere and allowed limited O_2_-dependent UQ biosynthesis in cells grown anaerobically. Thus, modifications (detailed below) were adopted in additional experiments conducted under strict anaerobic conditions.

### Culture under strict anaerobic conditions and cells quenching

LB medium was supplemented with 100 mg/L L-cysteine (adjusted to pH 6 with NaOH) and 2.5 mg/L resazurin. The medium was distributed in Hungate tubes and was deoxygenated by argon (O_2_<0.1 ppm) bubbling for 45 min at 60°C. The resazurin was initially purple, then quickly turned to pink, and eventually became colorless. The Hungate tubes were sealed and autoclaved. Two sequential precultures were performed in order to dilute the UQ present in the initial aerobic inoculum. The first preculture was performed overnight and used Eppendorf tubes filled to the top and inoculated with cells grown aerobically on LB agar. The second preculture was performed for 8 hours in Hungate tubes and was used to inocculate Hungate tubes that were subsequently incubated overnight at 37°C. Disposable syringes (1 mL) and needles were flushed 5 times with argon prior to inoculating 50 µL of preculture through the septum of the Hungate tubes. The resazurin remained colorless at all steps of the culture, indicating that the medium in the Hungate tubes was strictly anaerobic. At the end of the culture, the Hungate tubes were cooled down on ice for 45 min and 2 mL medium was sampled through the septum with argon-flushed syringes (2 mL) fitted with needles. The cells were immediately quenched by transfer to −20°C precooled glass tubes containing 6 mL methanol, 0.5 mL glass beads (0.5 mm diameter) and 20 mM KCl. The tubes were homogenized by vortex for 30 seconds and kept at −20°C prior to quinone extraction. In parallel, we also centrifuged 2 mL of culture from the Hungate tubes in order to determine the weight of the cells and normalize the UQ content of the quenched cells that was subsequently measured.

For the experiments conducted under strict anaerobic conditions (Figure 2H, 2I), we used LB medium instead of MSGN, since nitrite - produced during the anaerobic respiration of nitrate in MSGN medium - is able to oxidize resazurin [69].

### Lipid extraction and quinone analysis

Quinone extraction from cell pellets was performed as previously described [17].

Quinone extraction from cells quenched in methanol was slightly adapted from [9]. Briefly, 4 µL of a 10 µM UQ_10_ solution was added as internal standard to the cells-methanol mixture. Then 4 mL petroleum ether (boiling range 40–60 °C) was added, the tubes were vortexed for 30 sec and the phases were separated by centrifugation 1 min, 600 rpm. The upper petroleum ether layer was transferred to a fresh glass tube. Petroleum ether (4 mL) was added to the glass beads and methanol-containing tube, and the extraction was repeated. The petroleum ether layers were combined and dried under nitrogen.

The dried lipid extracts were resuspended in 100 µL ethanol and a volume corresponding to 1 mg of cells wet weight was analyzed by HPLC-electrochemical detection-mass spectrometry (ECD-MS) with a BetaBasic-18 column at a flow rate of 1 mL/min with mobile phases composed of methanol, ethanol, acetonitrile and a mix of 90% isopropanol, 10% ammonium acetate (1 M), 0.1% TFA: mobile phase 1 (50% methanol, 40% ethanol and 10% mix), mobile phase 2 (40% acetonitrile, 40% ethanol, 20% mix). Mobile phase 1 was used in MS detection on a MSQ spectrometer (Thermo Scientific) with electrospray ionization in positive mode (probe temperature 400°C, cone voltage 80V). Single ion monitoring (SIM) detected the following compounds: OPP (M+NH_4_^+^), m/z 656.0-656.8, 5-10 min, scan time 0.2 s; DMQ_8_ (M+Na_4_^+^), m/z 719-720, 6-10 min, scan time 0.2 s; ^13^C_6_ –DMQ_8_ (M+Na^+^), m/z 725-726, 6-10 min, scan time 0.2 s; UQ_8_ (M+ _4_^+^), m/z 744-745, 6-10 min, scan time 0.2 s; UQ_8_ (M+Na^+^), m/z 749-750, 6-10 min, scan time 0.2 s; ^13^C_6_-UQ_8_ (M+Na^+^), m/z 755.0-756, 6-10 min, scan time 0.2 s; UQ_10_ (M+NH_4_^+^), m/z 880.2-881.2, 10-17 min, scan time 0.2 s. MS spectra were recorded between m/z 600 and 900 with a scan time of 0.3 s. UV detection at 247 nm was used to quantify DMK_8_ and MK_8_. ECD, MS and UV peak areas were corrected for sample loss during extraction on the basis of the recovery of the UQ_10_ internal standard and were then normalized to cell’s wet weight. The peak of UQ_8_ obtained with electrochemical detection was quantified with a standard curve of UQ_10_ [17]. The absolute quantification of UQ_8_ based on the m/z= 744.6 signal at 8 min (Figure 2H, I) was performed with a standard curve of UQ_8_ ranging from 0.5 to 150 pmoles UQ_8_ (the detection limit was around 0.1 pmole).

### Anaerobic ^13^C_6_-UQ_8_ biosynthesis activity assay

Δ*ubi*C Δ*ubi*F cells containing or not the additional Δ*ubiU* or Δ*ubiV* deletions were grown overnight in MS medium supplemented with 0.2% glucose. This preculture was used to inoculate at OD_600_=0.1, 100 mL of fresh medium supplemented with 10 µM ^13^C_7_-4HB. The culture was grown at 37°C, 180 rpm until OD_600_=1, at which point 100 µM 4HB was added. The cells were pelleted by centrifugation at 3200 g, 4°C, 10 min and suspended in 100 mL MSGN medium. A 10 mL aliquot was taken for quinone extraction (aerobiosis, Figure 4B-D) and the rest of the culture was placed at 37°C in an anaerobic bottle with a two-port cap fitted with plastic tubing used to inject argon (O_2_<0.1 ppm) throughout the experiment in order to create and maintain anaerobiosis. After 5 min bubbling, a 10 mL sample was taken corresponding to 0 min anaerobiosis, then samples were taken every 30 minutes and analyzed for quinone content.

### Overexpression and purification of proteins

#### Overexpression and purification E. coli wild-type UbiV and variants

The pET-22b(+) plasmid, encoding wild-type UbiV or variants, were co-transformed with pGro7 plasmid (Takara Bio Inc.) into *E. coli* BL21 (DE3) competent cells. Single colonies obtained from transformation were grown overnight at 37°C in LB medium supplemented with ampicillin (50 µg/mL) and chloramphenicol (12.5 µg/mL). 10 mL of preculture was used to inoculate 1 L of LB medium with the same antibiotics, and the bacteria were cultured further at 37 °C with shaking (200 rpm). At an OD_600_ of 1.2, D-arabinose was added to the cultures at a final concentration of 2 mg/mL. At an OD_600_ of 1.8, the culture was cooled in an ice-water bath, and isopropyl 1-thio-β-D-galactopyranoside (IPTG) was added at a final concentration of 0.1 mM. Cells were then allowed to grow further at 16 °C overnight. All subsequent operations were carried out at 4°C. Cells were harvested in an Avanti^®^ J-26XP High-Performance centrifuge from Beckman Coulter with a JLA-8.1000 rotor at 5,000 × g for 10 minutes. The cell pellets were resuspended in 5 volumes of buffer A (50 mM Tris-HCl, pH 8.5, 150 mM NaCl, 15% (v/v) glycerol, 1 mM DTT) containing Complete™ Protease Inhibitor Cocktail (one tablet per 50 mL) (Roche) and disrupted by sonication (Branson Digital Sonifier, amplitude 40% for 10 min). Cells debris were removed by ultracentrifugation in an Optima™ XPN-80 ultracentrifuge from Beckman Coulter with a 50.2 Ti rotor at 35,000 x g for 60 min. The resulting supernatant was loaded onto a HisTrap FF Crude column (GE Healthcare) pre-equilibrated with buffer A. The column was washed with 10 column volumes of buffer B (50 mM Tris-HCl pH 8.5, 150 mM NaCl, 15% (v/v) glycerol, 1 mM DTT, 10 mM imidazole) to remove non-specifically bound *E. coli* proteins then eluted with a linear gradient of 10 column volumes of buffer B containing 500 mM imidazole. Fractions containing WT UbiV or variants were pooled and phenylmethylsulfonyl fluoride was added at a final concentration of 1mM. The proteins were then loaded on a HiLoad 16/600 Superdex 75 pg (GE Healthcare) pre-equilibrated in buffer C (50 mM Tris-HCl pH 8.5, 25 mM NaCl, 15% (v/v) glycerol, 1 mM DTT). The purified proteins were concentrated to 30-40 mg/mL using Amicon concentrators (30-kDa cutoff; Millipore), aliquoted, frozen in liquid nitrogen and stored at −80 °C. Overall, a high yield of 150 mg UbiV/L culture was obtained.

#### Overexpression and purification of UbiU/V complex and variants

The overexpression of wild-type UbiU/V complex or variants in *E. coli* BL21 (DE3) competent cells were performed following the same protocol as for UbiV. The expression of these proteins was induced by addition of IPTG to a final concentration of 0.05 mM. Wild-type UbiU/V complex or variants were purified with the same procedure as for UbiV, with the exception that the proteins were loaded on the HiLoad 16/600 Superdex 75 pg with buffer A.

### [Fe-S] cluster reconstitution

The [Fe-S] cluster(s) reconstitution of holo-UbiV and holo-UbiU/V were conducted under anaerobic conditions in an Mbraun LabStar glove box containing less than 0.5 ppm O_2_. Classically, a solution containing 100 µM of as-purified UbiV or UbiU/V complex, was treated with 5 mM DTT for 15 min at 20°C and then incubated for 1 hour with a 5-fold molar excess of both ferrous ammonium sulfate and L-cysteine. The reaction was initiated by the addition of a catalytic amount of the E.coli cysteine desulfurase CsdA (1-2% molar equivalent) and monitored by UV-visible absorption spectroscopy. The holo-UbiV or holo-UbiU/V complexes were then loaded onto a Superdex 75 Increase 10/300 GL column (GE Healthcare) pre-equilibrated with buffer C or A, respectively, to remove all excess of iron and L-cysteine. The fractions containing the holo-proteins were pooled and concentrated to 20-30mg/mL on a Vivaspin concentrator (30-kDa cutoff).

### Quantification Methods

Protein concentrations were determined using the method of Bradford (Bio-Rad) with bovine serum albumin as the standard. The iron and acid-labile sulfide were determined according to the method of Fish [70] and Beinert [71], respectively.

### UV-Vis spectroscopy

UV-Visible spectra were recorded in 1 cm optic path quartz cuvette under aerobic conditions on a Cary 100 UV-Vis spectrophotometer (Agilent) and under anaerobic conditions in a glove box on a XL-100 Uvikon spectrophotometer equipped with optical fibers.

### EPR spectroscopy

EPR spectra of frozen solutions were recorded on a Bruker Continuous-Waves (CW) X-Band ELEXSYS E500 spectrometer operating at 9.39 GHz, equipped with an SHQE cavity cooled by an helium flow cryostat ESR 900 Oxford Instruments under non-saturating conditions and using the following parameters: a microwave power in the range 2 to 10 mW and a modulation of the magnetic field at 100 kHz with a modulation amplitude of 0.6 mT. Holo UbiV or holo UbiU-UbiV complex were treated with 10-fold molar excess of dithionite to reduce the Fe-S cluster. Each solution was introduced into EPR quartz tubes in a glove box and frozen with liquid nitrogen before the EPR measurements.

### Genome datasets

The protein sequences from 5750 complete genomes (“extended dataset”) were downloaded from the NCBI Refseq database (bacteria and archaea, last accessed in November 2016, Table S2A). A representative set of complete genomes from a monophyletic group of bacteria that potentially harbor the *ubi*quinone biosynthesis pathway was also created: “Reference” and “Representative” genomes were downloaded from the NCBI Refseq database for 204 Alphaproteobacteria, 103 Betaproteobacteria, 303 Gammaproteobacteria (last accessed in November 2018). In addition to these 610 genomes, the genome of *Phaeospirillum fulvum* (99.5% estimated completeness according to CheckM, http://gtdb.ecogenomic.org/genomes?gid=GCF_900108475.1) was included (Table S2B).

### HMM protein profiles creation

An initial set of protein sequences (“curated set”) was retrieved from genomes manually and from a publication [72], to cover the diversity of UQ-producing organisms. The curated set included 48 pairs of YhbU and YhbV from 10 alpha, 19 beta and 19 gamma-proteobacteria, 17 sequences for YhbT, 64, 181, 69 and 189 sequences for UbiA, MenA, UbiG and UbiE respectively (Table S4). Then, for each gene family these sequences were aligned with Mafft (v7.313, “linsi”) [73], and each alignment trimmed at its N-ter and C-ter extremities based on the filtering results of BMGE (BLOSUM 30) [74]. The core of the alignments were kept as is, and Hidden Markov Model (HMM) profiles were created directly from the trimmed alignments using the Hmmbuild program (Hmmer suite, version 3.1b2) [75].

To ensure a more sensitive search, and good delineation between homologs, phylogenetic curation was used, as YhbU and YhbV are known to be part of the larger U32 protease gene family [43]. A search using the YhbU and YhbV HMM profiles was performed with Hmmsearch (Hmmer suite) on the extended 5750 genomes dataset, and sequences with an i-evalue (“independent e-value”) lower than 10E-20 and a coverage of the profiles higher than 90% were selected. The 4212 sequences obtained were de-replicated using Uclust from the Usearch program suite (80% identity level) [76]. A phylogenetic tree was built by maximum-likelihood with the IQ-Tree program (best evolutionary model) based on the alignment (Mafft: “linsi”, BMGE with BLOSUM30) of the 480 selected sequences including all curated YhbU and YhbV sequences [73,74,77]. YhbU and YhbV proteobacterial sequences formed two separate monophyletic groups, giving credit to our curated set (100% and 80% UF-Boot support respectively). The other sequences that formed a large monophyletic group of bacterial sequences were categorized as “U32 proteases” (98% UF-Boot, https://doi.org/10.6084/m9.figshare.7800614.v1). The 98 sequences from this U32 proteases group (Table S4) were used to re-create an HMM profile as described above, and served as an outgroup for the profile search.

For YhbT, the first profile obtained from the curated set of sequences was used altogether with the YhbU, YhbV and U32 proteases profiles for a search in the 611 proteobacteria genomes dataset. A second profile was created from YhbT sequences (YhbT2, Table S4) that were co-localizing with YhbU and YhbV hits (10E-20 i-evalue and 80% profile coverage). The two YhbT profiles matched complementary sets of sequences and therefore were both used for annotating YhbT in genomes.

A similar approach was taken in order to identify the six known aerobic hydroxylases. 51 Coq7, 73 UbiF, 80 UbiH, 58 UbiI, 24 UbiL and 32 UbiM sequences were extracted manually and from publications [72] [19](Table S4, “version 1”) to serve as a reference, annotated set of sequences. Profiles were created as described above. To ensure their specificity, we ran the HMM profiles against our 5570 genomes dataset and selected the sequences that had an i-evalue lower than 10E-20 and a coverage of the profiles higher than 90%. We built two phylogenetic trees as described above: one for Coq7, and another one for UbiFHILM, which are known to be part of the large FMO protein family [19]. In the latter case, we de-replicated the 1619 sequences obtained for the FMO protein family before performing the alignment, alignment filtering, and tree reconstruction steps (using Uclust at the 60% identity level). The Coq7 tree obtained showed our reference Coq7 sequences covered the whole diversity of retrieved sequences, suggesting that they all could be *bona fide* Coq7 (https://doi.org/10.6084/m9.figshare.7800680). The FMO tree showed a monophyletic group containing all reference FMO *ubi*quinone hydroxylases, forming sub-groups for the different homologs (UbiFHILM) in Proteobacteria (https://doi.org/10.6084/m9.figshare.7800620). Further, a large set of sequences formed an outgroup consisting of sequences from various clades of bacteria, a lot being found outside of Proteobacteria, robustly separated from the *ubi*quinone hydroxylases. We split this large clade into four sub-trees, and extracted the corresponding sequences to obtain four new HMM profiles (as described above), to be used for precise discrimination between *ubi*quinone hydroxylases and other members of the FMO family (“AlloFMO_1” to “AlloFMO_4” in Table S4, “version 2”). FMO *ubi*quinone hydroxylases sub-trees were also used to re-design improved HMM profiles for UbiFHILM (36, 168, 198, 139, and 65 sequences respectively, see Table S4 “version 2”).

### Evaluation of genomic distributions with HMMER and MacSyFinder

Two MacSyFinder models were created to i) search sequences of interest in genomes using Hmmer, and ii) investigate their genetic architecture [78]. A first model was created to focus only on YhbTUV-related genes. In this model, the YhbTUV components were defined as “mandatory”, and U32 protease as “accessory”. A second, more comprehensive model “Ubi+YhbTUV” was designed to list the families corresponding to the 9 profiles obtained (UbiA, MenA, UbiE, UbiG, 2 YhbT, YhbU, YhbV and U32 proteases). For both models, the two YhbT profiles were set as “exchangeable”. The parameter “inter_gene_max_space” was set to 5 in the “YhbTUV” model, and 10 in the “Ubi+YhbTUV” model. MacSyFinder was set to run HMMER with the options “--i-evalue-select 10E-20” and “--coverage-profile 0.8”. Independently of their genetic context, sequences corresponding to selected HMMER hits were listed for all profiles in all genomes analyzed in order to establish the genomic distribution for each of the protein families of interest. When several profiles matched a sequence, only the best hit (best i-evalue) was considered.

### U32 proteases sequence analysis

We retrieved from the UniProt-KB database 3460 protein sequences of U32 proteases that were categorized in 12 families by Kimura *et al*. [43], and created a FASTA for each of these families. 50 sequences from different families could not be retrieved as they had been deleted from the Uniprot-KB database (46 “obsolete”) or were not found based on the published accession numbers. As the “RlhA1” and “RlhA2” families mostly corresponded to two domains from the same protein sequences that had been split, we put whole sequences all together into a single fasta file for sequence analysis of the overall “RlhA” family. For each of the 10 family, sequences were de-replicated at the 80% identity level with Uclust, in order to limit any potential taxonomic sampling bias, and sequences were aligned (Mafft, linsi). The alignments were visualized in Jalview [79] and used to create the logo sequences. Images of alignments were created using the ESPript webserver (http://espript.ibcp.fr/ESPript/ESPript/) [80].

## ACKNOWLEDGEMENTS

This work was supported by the Agence Nationale de la Recherche (ANR), ANR Blanc (An)aeroUbi ANR-15-CE11-0001-02 to FP. We thank Amélie Amblard for technical assistance, Louis Givelet for preliminary bioinformatic analyses, Barbara Schoepp-Cothenet for providing accession numbers to sequences of UbiA, -G, -E and for critical reading of the manuscript, and the GEM team at TIMC for discussions and suggestions. CDTV, ML and MF acknowledge support from the French National Research Agency (Labex program DYNAMO, ANR-11-LABX-0011). CC was funded by the Grenoble Alpes Data Institute, supported by the French National Research Agency under the “Investissements d’avenir” program (ANR-15-IDEX-02). We thank the National Bioresource Project, National Institute of Genetics for providing ME strains from the medium and large deletions *E. coli* collection.

## SUPPLEMENTARY FIGURES LEGENDS

**Figure S1:** A) HPLC-ECD analyses (mobile phase 2) of lipid extracts from 1 mg of WT, Δ*ubiT* and Δ*ubiG* cells grown in LB medium under anaerobic conditions (UQ_10_ used as standard). B) Genetic context of the *ubiT,U,V* genes in M. marinus (WP_011715033.1). The scheme was drawn from a figure obtained using the GeneSpy program [81].

**Figure S2:** Multiple sequence alignment of UbiT from representative proteobacteria. Sequences were aligned using Mafft (linsi), and the output generated using Jalview and Inkscape. Hydrophobic residues are colored in blue. The position of the SCP2 domain (PF02036) is indicated by a red box. Genbank accession numbers: *Vibrio cholerae*, NP_230303.1, *Pseudomonas aeruginosa*, NP_252600.1, *Escherichia coli*, NP_312065.1, *Yersinia pestis*, YP_002348366.1, *Ralstonia solanacearum*, WP_011004260.1, *Dechloromonas aromatica*, WP_011285876.1, *Thiobacillus denitrificans*, WP_011312695.1, *Rhodopseudomonas palustris*, WP_011666115.1, *Halorhodospira halophila*, WP_011813831.1, *Rhodobacter sphaeroides*, WP_011909347.1, *Aeromonas salmonicida*, WP_005310396.1, *Paracoccus denitrificans*, WP_011750456.1, *Neisseria sicca*, WP_080614297.1, *Phaeospirillum fulvum*, WP_074767212.1.

**Figure S3:** Multiple sequence alignment of UbiU from representative proteobacteria. Sequences were aligned using Mafft (linsi), and the output generated using ESPript and Inkscape. The four conserved cysteines (C169, C176, C193 and C232) involved in iron-sulfur binding are indicated by black columns. The position of the domain U32 protease PF01136 is indicated by a green box. GenBank accession numbers: *Vibrio cholerae*, NP_230301.1, *Pseudomonas aeruginosa*, NP_252602.1, *Escherichia coli*, NP_312066.1, *Yersinia pestis*, YP_002348368.1, *Ralstonia solanacearum*, WP_011004262.1, *Dechloromonas aromatica*, WP_011285878.1, *Thiobacillus denitrificans*, WP_011312698.1, *Rhodopseudomonas palustris*, WP_011666114.1, *Halorhodospira halophila*, WP_011813833.1, *Rhodobacter sphaeroides*, WP_011909346.1, *Aeromonas salmonicida*, WP_005310394.1, *Paracoccus denitrificans*, WP_041530457.1, *Neisseria sicca*, WP_049226489.1, *Phaeospirillum fulvum*, WP_074767209.1.

**Figure S4:** Multiple sequence alignment of UbiV from representative proteobacteria. Sequences were aligned using Mafft (linsi), and the output generated using ESPript and Inkscape. The four conserved cysteines (C39, C180, C193 and C197 - positions in *E. coli*) involved in iron-sulfur binding are indicated by black columns. The position of the domain U32 protease PF01136 is indicated by a green box. GenBank accession numbers: *Vibrio cholerae*, NP_230300.2, *Pseudomonas aeruginosa*, NP_252601.1, *Escherichia coli*, NP_312067.2, Yersinia pestis, YP_002348369.1, *Ralstonia solanacearum*, WP_011004261.1, *Dechloromonas aromatica*, WP_011285877.1, *Thiobacillus denitrificans*, WP_011312697.1, *Rhodopseudomonas palustris*, WP_011666113.1, *Halorhodospira halophila*, WP_011813832.1, *Rhodobacter sphaeroides*, WP_011909345.1, *Aeromonas salmonicida*, WP_005310392.1, *Paracoccus denitrificans*, WP_011750458.1, *Neisseria sicca*, WP_080614296.1, *Phaeospirillum fulvum*, WP_074767477.1.

**Figure S5:** A) Gel filtration chromatogram of aerobically purified UbiV on a Superdex 75 Increase 10/300 GL column; Inset: Coomassie staining SDS-PAGE of aerobically purified UbiV. B) Calibration curve, standard proteins in rhombi, UbiV in circle. C) UV-visible absorption spectra of 41 µM holo-UbiV under anaerobiosis (solid line) and after 20 minutes exposure to air (dotted line). D) Comparative UV-visible absorption spectra of aerobically purified wild-type (black) and different Cys-to-Ala mutants of UbiV (C193A C197A in blue; C39A C193A C197A in green; C180A C193A C197A in pink; C39A C180A C193A C197A in brown). Inset: Coomassie staining SDS-PAGE shows the purifications of UbiV and variants. Lanes are: M, molecular mass marker; 1, wild-type UbiV; 2, UbiV C193A C197A; 3, UbiV C39A C193A C197A; 4, UbiV C180A C193A C197A; 5, UbiV C39A C180A C193A C197. All proteins were in 50 mM Tris-HCl, pH 8.5, 25 mM NaCl, 15% glycerol, 1 mM DTT (A-D).

**Figure S6:** A) Gel filtration chromatogram of aerobically purified UbiU-UbiV on a Superdex 75 Increase 10/300 GL column; Inset: Coomassie staining SDS-PAGE of aerobically purified UbiU-UbiV complex. B) Calibration curve, standard proteins in rhombi, UbiU-UbiV in circle. C) Coomassie staining SDS-PAGE shows the purifications of UbiU-UbiV wild-type and variants proteins: Lanes are: M, molecular mass marker; 1, wild-type UbiU-UbiV complex; 2, UbiU C169A C176A - UbiV wild-type; 3, UbiU C193A C232A - UbiV wild-type. All proteins were in buffer 50 mM Tris-HCl, pH 8.5, 150 mM NaCl, 15% glycerol, 1 mM DTT (A-D).

**Table S1:** UQ levels in strains from the medium- and large-deletion collections

**Table S2:** Occurrence of genes from the UQ pathways in bacterial genomes.

**Table S3:** List of oligonucleotides and strains used in this study

**Table S4:** References of the protein sequences used to build the HMM profiles.

## REFERENCES

1. Schoepp-Cothenet B, van Lis R, Atteia A, Baymann F, Capowiez L, Ducluzeau AL, et al. On the universal core of bioenergetics. Biochim Biophys Acta. 2013; 1827: 79–93.

2. Marteyn B, Scorza FB, Sansonetti PJ, Tang C. Breathing life into pathogens: the influence of oxygen on bacterial virulence and host responses in the gastrointestinal tract. Cell Microbiol. 2011; 13: 171–176.

3. Wallace N, Zani A, Abrams E, Sun Y (2016) The Impact of Oxygen on Bacterial Enteric Pathogens. In: Sariaslani S, Gadd GM, editors. Advances in Applied Microbiology, Vol 95. San Diego: Elsevier Academic Press Inc. pp. 179–204.

4. Bettenbrock K, Bai H, Ederer M, Green J, Hellingwerf KJ, Holcombe M, et al. Towards a Systems Level Understanding of the Oxygen Response of Escherichia coli. Advances in Microbial Systems Biology. 2014; 64: 65–114.

5. Morris RL, Schmidt TM. Shallow breathing: bacterial life at low O-2. Nature Reviews Microbiology. 2013; 11: 205–212.

6. Marreiros BC, Calisto F, Castro PJ, Duarte AM, Sena FV, Silva AF, et al. Exploring membrane respiratory chains. Biochimica Et Biophysica Acta-Bioenergetics. 2016; 1857: 1039–1067.

7. Lecomte SM, Achouak W, Abrouk D, Heulin T, Nesme X, Haichar FE. Diversifying Anaerobic Respiration Strategies to Compete in the Rhizosphere. Frontiers in Environmental Science. 2018; 6: 16.

8. Nowicka B, Kruk J. Occurrence, biosynthesis and function of isoprenoid quinones. Biochim Biophys Acta. 2010; 1797: 1587–1605.

9. Sharma P, Teixeira de Mattos MJ, Hellingwerf KJ, Bekker M. On the function of the various quinone species in Escherichia coli. FEBS J. 2012; 279: 3364–3373.

10. Shestopalov AI, Bogachev AV, Murtazina RA, Viryasov MB, Skulachev VP. Aeration-dependent changes in composition of the quinone pool in Escherichia coli - Evidence of post-transcriptional regulation of the quinone biosynthesis. Febs Letters. 1997; 404: 272–274.

11. Nitzschke A, Bettenbrock K. All three quinone species play distinct roles in ensuring optimal growth under aerobic and fermentative conditions in E. coli K12. PLoS One. 2018; 13: e0194699.

12. Aussel L, Pierrel F, Loiseau L, Lombard M, Fontecave M, Barras F. Biosynthesis and physiology of coenzyme Q in bacteria. Biochim Biophys Acta. 2014; 1837: 1004–1011.

13. Reidenbach AG, Kemmerer ZA, Aydin D, Jochem A, McDevitt MT, Hutchins PD, et al. Conserved Lipid and Small-Molecule Modulation of COQ8 Reveals Regulation of the Ancient Kinase-like UbiB Family. Cell Chem Biol. 2018; 25: 154–165 e111.

14. Loiseau L, Fyfe C, Aussel L, Hajj Chehade M, Hernandez SB, Faivre B, et al. The UbiK protein is an accessory factor necessary for bacterial ubiquinone (UQ) biosynthesis and forms a complex with the UQ biogenesis factor UbiJ. J Biol Chem. 2017; 292: 11937–11950.

15. Aussel L, Loiseau L, Hajj Chehade M, Pocachard B, Fontecave M, Pierrel F, et al. ubiJ, a New Gene Required for Aerobic Growth and Proliferation in Macrophage, Is Involved in Coenzyme Q Biosynthesis in Escherichia coli and Salmonella enterica Serovar Typhimurium. J Bacteriol. 2014; 196: 70–79.

16. Hajj Chehade M, Pelosi L, Fyfe CD, Loiseau L, Rascalou B, Brugiere S, et al. A Soluble Metabolon Synthesizes the Isoprenoid Lipid Ubiquinone. Cell Chem Biol. 2019; 26: 482–492 e487.

17. Hajj Chehade M, Loiseau L, Lombard M, Pecqueur L, Ismail A, Smadja M, et al. ubiI, a New Gene in Escherichia coli Coenzyme Q Biosynthesis, Is Involved in Aerobic C5-hydroxylation. J Biol Chem. 2013; 288: 20085–20092.

18. Alexander K, Young IG. Three hydroxylations incorporating molecular oxygen in the aerobic biosynthesis of ubiquinone in Escherichia coli. Biochemistry. 1978; 17: 4745–4750.

19. Pelosi L, Ducluzeau AL, Loiseau L, Barras F, Schneider D, Junier I, et al. Evolution of Ubiquinone Biosynthesis: Multiple Proteobacterial Enzymes with Various Regioselectivities To Catalyze Three Contiguous Aromatic Hydroxylation Reactions. mSystems. 2016; 1: e00091–00016.

20. Alexander K, Young IG. Alternative hydroxylases for the aerobic and anaerobic biosynthesis of ubiquinone in Escherichia coli. Biochemistry. 1978; 17: 4750–4755.

21. Cox GB, Gibson F. Biosynthesis of Vitamin K and Ubiquinone. Relation to the Shikimic Acid Pathway in Escherichia Coli. Biochim Biophys Acta. 1964; 93: 204–206.

22. Siebert M, Severin K, Heide L. Formation of 4-hydroxybenzoate in Escherichia coli: characterization of the ubiC gene and its encoded enzyme chorismate pyruvate-lyase. Microbiology. 1994; 140 (Pt 4): 897–904.

23. Payet LA, Leroux M, Willison JC, Kihara A, Pelosi L, Pierrel F. Mechanistic Details of Early Steps in Coenzyme Q Biosynthesis Pathway in Yeast. Cell Chem Biol. 2016; 23: 1241–1250.

24. Kwon O, Kotsakis A, Meganathan R. Ubiquinone (coenzyme Q) biosynthesis in Escherichia coli: identification of the ubiF gene. FEMS Microbiol Lett. 2000; 186: 157–161.

25. Hashimoto M, Ichimura T, Mizoguchi H, Tanaka K, Fujimitsu K, Keyamura K, et al. Cell size and nucleoid organization of engineered Escherichia coli cells with a reduced genome. Mol Microbiol. 2005; 55: 137–149.

26. Mizoguchi H, Sawano Y, Kato J, Mori H. Superpositioning of deletions promotes growth of Escherichia coli with a reduced genome. DNA Res. 2008; 15: 277–284.

27. Baba T, Ara T, Hasegawa M, Takai Y, Okumura Y, Baba M, et al. Construction of Escherichia coli K-12 in-frame, single-gene knockout mutants: the Keio collection. Mol Syst Biol. 2006; 2: 2006 0008.

28. Esposti MD. A journey across genomes uncovers the origin of ubiquinone in cyanobacteria. Genome Biol Evol. 2017.

29. Hiraishi A, Hoshino Y, Kitamura H. Isoprenoid quinone composition in the classification of rhodospirillaceae. Journal of General and Applied Microbiology. 1984; 30: 197–210.

30. Imhoff JF, Petri R, Suling J. Reclassification of species of the spiral-shaped phototrophic purple non-sulfur bacteria of the alpha-Proteobacteria: description of the new genera Phaeospirillum gen. nov., Rhodovibrio gen. nov., Rhodothalassium gen. nov. and Roseospira gen. nov. as well as transfer of Rhodospirillum fulvum to Phaeospirillum fulvum comb. nov., of Rhodospirillum molischianum to Phaeospirillum molischianum comb. nov., of Rhodospirillum salinarum to Rhodovibrio salinarum comb. nov., of Rhodospirillum sodomense to Rhodovibrio sodomensis comb. nov., of Rhodospirillum salexigens to Rhodothalassium salexigens comb. nov, and of Rhodospirillum mediosalinum to Roseospira mediosalina comb. nov. International Journal of Systematic Bacteriology. 1998; 48: 793–798.

31. Bazylinski DA, Williams TJ, Lefevre CT, Berg RJ, Zhang CL, Bowser SS, et al. Magnetococcus marinus gen. nov., sp. nov., a marine, magnetotactic bacterium that represents a novel lineage (Magnetococcaceae fam. nov., Magnetococcales ord. nov.) at the base of the Alphaproteobacteria. Int J Syst Evol Microbiol. 2013; 63: 801–808.

32. Stewart CS, Duncan SH, Cave DR. Oxalobacter formigenes and its role in oxalate metabolism in the human gut. FEMS Microbiol Lett. 2004; 230: 1–7.

33. Burgardt NI, Gianotti AR, Ferreyra RG, Ermacora MR. A structural appraisal of sterol carrier protein 2. Biochimica Et Biophysica Acta-Proteins and Proteomics. 2017; 1865: 565–577.

34. Yano T, Yagi T, Sled VD, Ohnishi T. Expression and characterization of the 66-kilodalton (NQO3) iron-sulfur subunit of the proton-translocating NADH-quinone oxidoreductase of Paracoccus denitrificans. J Biol Chem. 1995; 270: 18264–18270.

35. Ollagnier S, Mulliez E, Gaillard J, Eliasson R, Fontecave M, Reichard P. The anaerobic Escherichia coli ribonucleotide reductase. Subunit structure and iron sulfur center. J Biol Chem. 1996; 271: 9410–9416.

36. Ollagnier de Choudens S, Barras F. Genetic, Biochemical, and Biophysical Methods for Studying FeS Proteins and Their Assembly. Methods Enzymol. 2017; 595: 1–32.

37. M NM Ollagnier-de-Choudens S, Sanakis Y, Abdel-Ghany SE, Rousset C, Ye H, et al. Characterization of Arabidopsis thaliana SufE2 and SufE3: functions in chloroplast iron-sulfur cluster assembly and Nad synthesis. J Biol Chem. 2007; 282: 18254–18264.

38. Ollagnier-De Choudens S, Sanakis Y, Hewitson KS, Roach P, Baldwin JE, Munck E, et al. Iron-sulfur center of biotin synthase and lipoate synthase. Biochemistry. 2000; 39: 4165–4173.

39. Ollagnier-de Choudens S, Loiseau L, Sanakis Y, Barras F, Fontecave M. Quinolinate synthetase, an iron-sulfur enzyme in NAD biosynthesis. FEBS Lett. 2005; 579: 3737–3743.

40. Hewitson KS, Ollagnier-de Choudens S, Sanakis Y, Shaw NM, Baldwin JE, Munck E, et al. The iron-sulfur center of biotin synthase: site-directed mutants. J Biol Inorg Chem. 2002; 7: 83–93.

41. Johnson DC, Dean DR, Smith AD, Johnson MK. Structure, function, and formation of biological iron-sulfur clusters. Annu Rev Biochem. 2005; 74: 247–281.

42. Fontecave M, Py B, Ollagnier de Choudens S, Barras F. From Iron and Cysteine to Iron-Sulfur Clusters: the Biogenesis Protein Machineries. EcoSal Plus. 2008; 3.

43. Kimura S, Sakai Y, Ishiguro K, Suzuki T. Biogenesis and iron-dependency of ribosomal RNA hydroxylation. Nucleic Acids Res. 2017; 45: 12974–12986.

44. Yen HW, Chiu CH. The influences of aerobic-dark and anaerobic-light cultivation on CoQ(10) production by Rhodobacter sphaeroides in the submerged fermenter. Enzyme and Microbial Technology. 2007; 41: 600–604.

45. Matsumura M, Kobayashi T, Aiba S. ANAEROBIC PRODUCTION OF UBIQUINONE-10 BY PARACOCCUS-DENITRIFICANS. European Journal of Applied Microbiology and Biotechnology. 1983; 17: 85–89.

46. Schoepp-Cothenet B, Lieutaud C, Baymann F, Vermeglio A, Friedrich T, Kramer DM, et al. Menaquinone as pool quinone in a purple bacterium. Proc Natl Acad Sci U S A. 2009; 106: 8549–8554.

47. Stolper DA, Revsbech NP, Canfield DE. Aerobic growth at nanomolar oxygen concentrations. Proceedings of the National Academy of Sciences of the United States of America. 2010; 107: 18755–18760.

48. D’Mello R, Hill S, Poole RK. The cytochrome bd quinol oxidase in Escherichia coli has an extremely high oxygen affinity and two oxygen-binding haems: implications for regulation of activity in vivo by oxygen inhibition. Microbiology. 1996; 142 (Pt 4): 755–763.

49. Rolfe MD, Ocone A, Stapleton MR, Hall S, Trotter EW, Poole RK, et al. Systems analysis of transcription factor activities in environments with stable and dynamic oxygen concentrations. Open Biol. 2012; 2: 120091.

50. Rolfe MD, Ter Beek A, Graham AI, Trotter EW, Asif HM, Sanguinetti G, et al. Transcript profiling and inference of Escherichia coli K-12 ArcA activity across the range of physiologically relevant oxygen concentrations. J Biol Chem. 2011; 286: 10147–10154.

51. Lee KM, Park Y, Bari W, Yoon MY, Go J, Kim SC, et al. Activation of Cholera Toxin Production by Anaerobic Respiration of Trimethylamine N-oxide in Vibrio cholerae. Journal of Biological Chemistry. 2012; 287: 11.

52. Zhao H, Li X, Johnson DE, Mobley HLT. Identification of protease and rpoN-associated genes of uropathogenic Proteus mirabilis by negative selection in a mouse model of ascending urinary tract infection. Microbiology-Uk. 1999; 145: 185–195.

53. Navais R, Mendez J, Perez-Pascual D, Cascales D, Guijarro JA. The yrpAB operon of Yersinia ruckeri encoding two putative U32 peptidases is involved in virulence and induced under microaerobic conditions. Virulence. 2014; 5: 619–624.

54. Groenewold MK, Massmig M, Hebecker S, Danne L, Magnowska Z, Nimtz M, et al. A phosphatidic acid binding protein is important for lipid homeostasis and adaptation to anaerobic biofilm conditions in Pseudomonas aeruginosa. Biochem J. 2018.

55. Filiatrault MJ, Picardo KF, Ngai H, Passador L, Iglewski BH. Identification of Pseudomonas aeruginosa genes involved in virulence and anaerobic growth. Infect Immun. 2006; 74: 4237–4245.

56. Schobert M, Jahn D. Anaerobic physiology of Pseudomonas aeruginosa in the cystic fibrosis lung. Int J Med Microbiol. 2010; 300: 549–556.

57. Line L, Alhede M, Kolpen M, Kuhl M, Ciofu O, Bjarnsholt T, et al. Physiological levels of nitrate support anoxic growth by denitrification of Pseudomonas aeruginosa at growth rates reported in cystic fibrosis lungs and sputum. Frontiers in Microbiology. 2014; 5: 11.

58. Behan RK, Lippard SJ. The aging-associated enzyme CLK-1 is a member of the carboxylate-bridged diiron family of proteins. Biochemistry. 2010; 49: 9679–9681.

59. Estellon J, Ollagnier de Choudens S, Smadja M, Fontecave M, Vandenbrouck Y. An integrative computational model for large-scale identification of metalloproteins in microbial genomes: a focus on iron-sulfur cluster proteins. Metallomics. 2014; 6: 1913–1930.

60. Rawlings ND, Alan J, Thomas PD, Huang XD, Bateman A, Finn RD. The MEROPS database of proteolytic enzymes, their substrates and inhibitors in 2017 and a comparison with peptidases in the PANTHER database. Nucleic Acids Research. 2018; 46: D624–D632.

61. Rouault TA. Mammalian iron-sulphur proteins: novel insights into biogenesis and function. Nature Reviews Molecular Cell Biology. 2015; 16: 45–55.

62. Lill R. Function and biogenesis of iron-sulphur proteins. Nature. 2009; 460: 831–838.

63. Roche B, Aussel L, Ezraty B, Mandin P, Py B, Barras F. Iron/sulfur proteins biogenesis in prokaryotes: formation, regulation and diversity. Biochim Biophys Acta. 2013; 1827: 455–469.

64. Hille R. Molybdenum-containing hydroxylases. Arch Biochem Biophys. 2005; 433: 107–116.

65. Datsenko KA, Wanner BL. One-step inactivation of chromosomal genes in Escherichia coli K-12 using PCR products. Proc Natl Acad Sci U S A. 2000; 97: 6640–6645.

66. Miller JH (1972) Experiments in Molecular Genetics; Laboratory CSH, editor.

67. Cherepanov PP, Wackernagel W. Gene disruption in Escherichia coli: TcR and KmR cassettes with the option of Flp-catalyzed excision of the antibiotic-resistance determinant. Gene. 1995; 158: 9–14.

68. Alberge F, Espinosa L, Seduk F, Sylvi L, Toci R, Walburger A, et al. Dynamic subcellular localization of a respiratory complex controls bacterial respiration. Elife. 2015; 4.

69. Jenneman GE, Montgomery AD, McInerney MJ. Method for detection of microorganisms that produce gaseous nitrogen oxides. Appl Environ Microbiol. 1986; 51: 776–780.

70. Fish WW. Rapid colorimetric micromethod for the quantitation of complexed iron in biological samples. Methods in Enzymology. 1988; 158: 357–364.

71. Beinert H. Micro methods for the quantitative determination of iron and copper in biological material. Methods in enzymology. 1978; 54: 435–445.

72. Ravcheev DA, Thiele I. Genomic Analysis of the Human Gut Microbiome Suggests Novel Enzymes Involved in Quinone Biosynthesis. Front Microbiol. 2016; 7: 128.

73. Katoh K, Standley DM. MAFFT multiple sequence alignment software version 7: improvements in performance and usability. Mol Biol Evol. 2013; 30: 772–780.

74. Criscuolo A, Gribaldo S. BMGE (Block Mapping and Gathering with Entropy): a new software for selection of phylogenetic informative regions from multiple sequence alignments. BMC Evol Biol. 2010; 10: 210.

75. Eddy SR. Accelerated Profile HMM Searches. PLoS Comput Biol. 2011; 7: e1002195.

76. Edgar RC. Search and clustering orders of magnitude faster than BLAST. Bioinformatics. 2010; 26: 2460–2461.

77. Nguyen LT, Schmidt HA, von Haeseler A, Minh BQ. IQ-TREE: a fast and effective stochastic algorithm for estimating maximum-likelihood phylogenies. Mol Biol Evol. 2015; 32: 268–274.

78. Abby SS, Neron B, Menager H, Touchon M, Rocha EP. MacSyFinder: a program to mine genomes for molecular systems with an application to CRISPR-Cas systems. PLoS One. 2014; 9: e110726.

79. Waterhouse AM, Procter JB, Martin DM, Clamp M, Barton GJ. Jalview Version 2--a multiple sequence alignment editor and analysis workbench. Bioinformatics. 2009; 25: 1189–1191.

80. Robert X, Gouet P. Deciphering key features in protein structures with the new ENDscript server. Nucleic Acids Res. 2014; 42: W320–324.

81. Garcia PS, Jauffrit F, Grangeasse C, Brochier-Armanet C. GeneSpy, a user-friendly and flexible genomic context visualizer. Bioinformatics (Oxford, England). 2019; 35: 329–331.

