## Supplementary figures and images for "Ubiquinone biosynthesis over the entire O_2_ range: characterization of a conserved, O_2_-independent pathway"

### Supplemental Figure 1

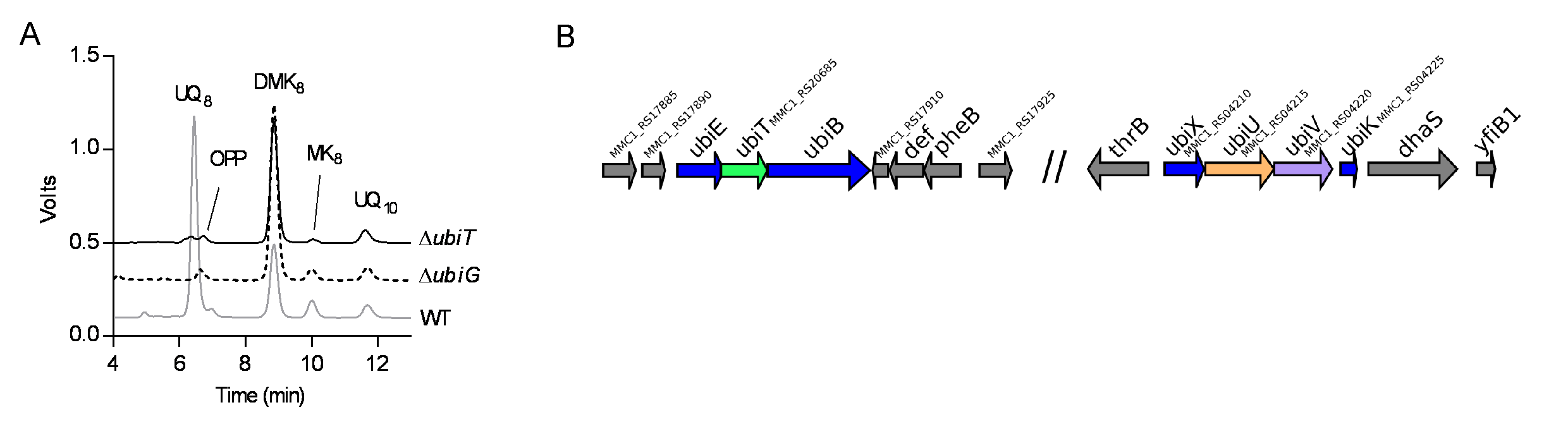

### Supplemental Figure 2

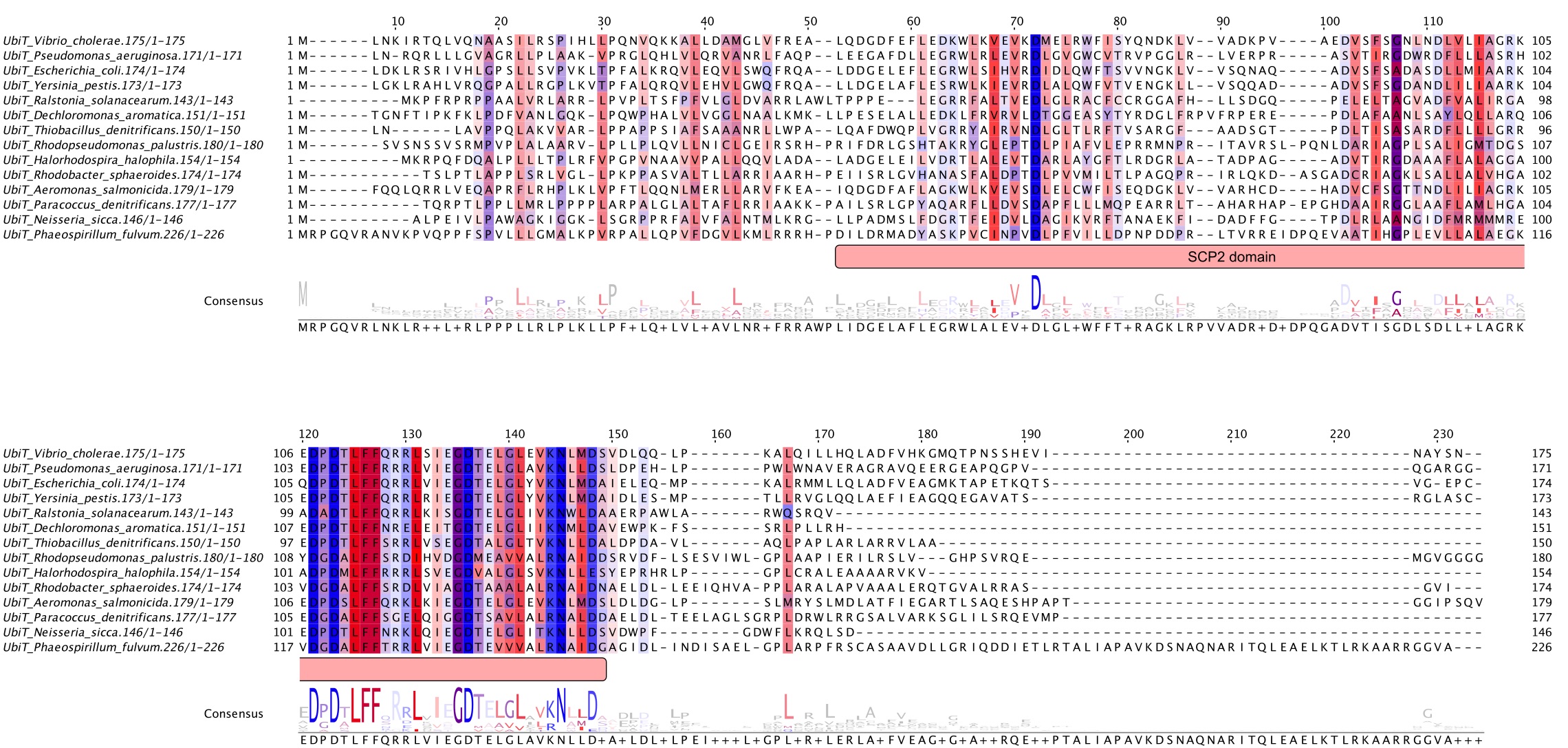

### Supplemental Figure 3

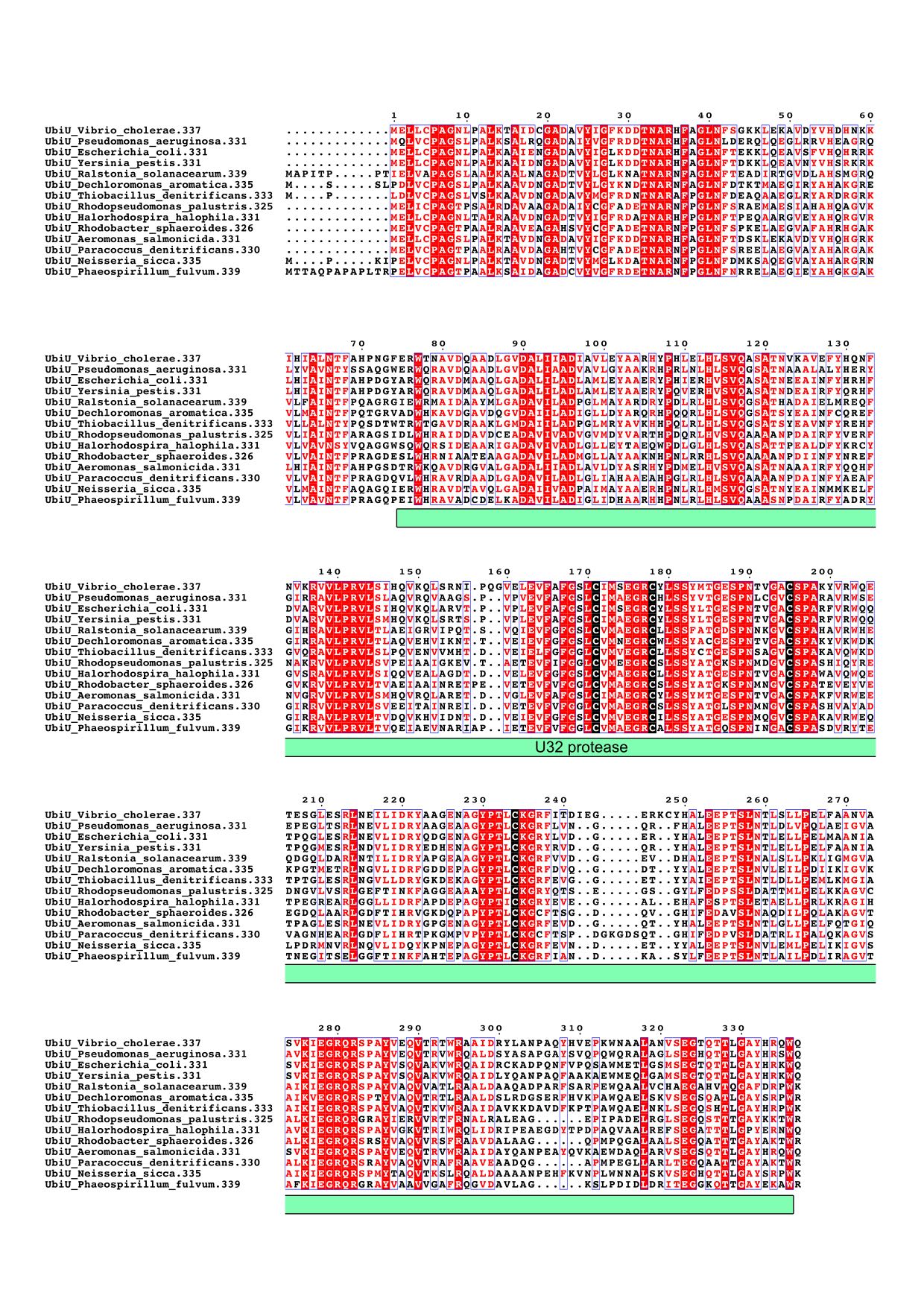

### Supplemental Figure 4

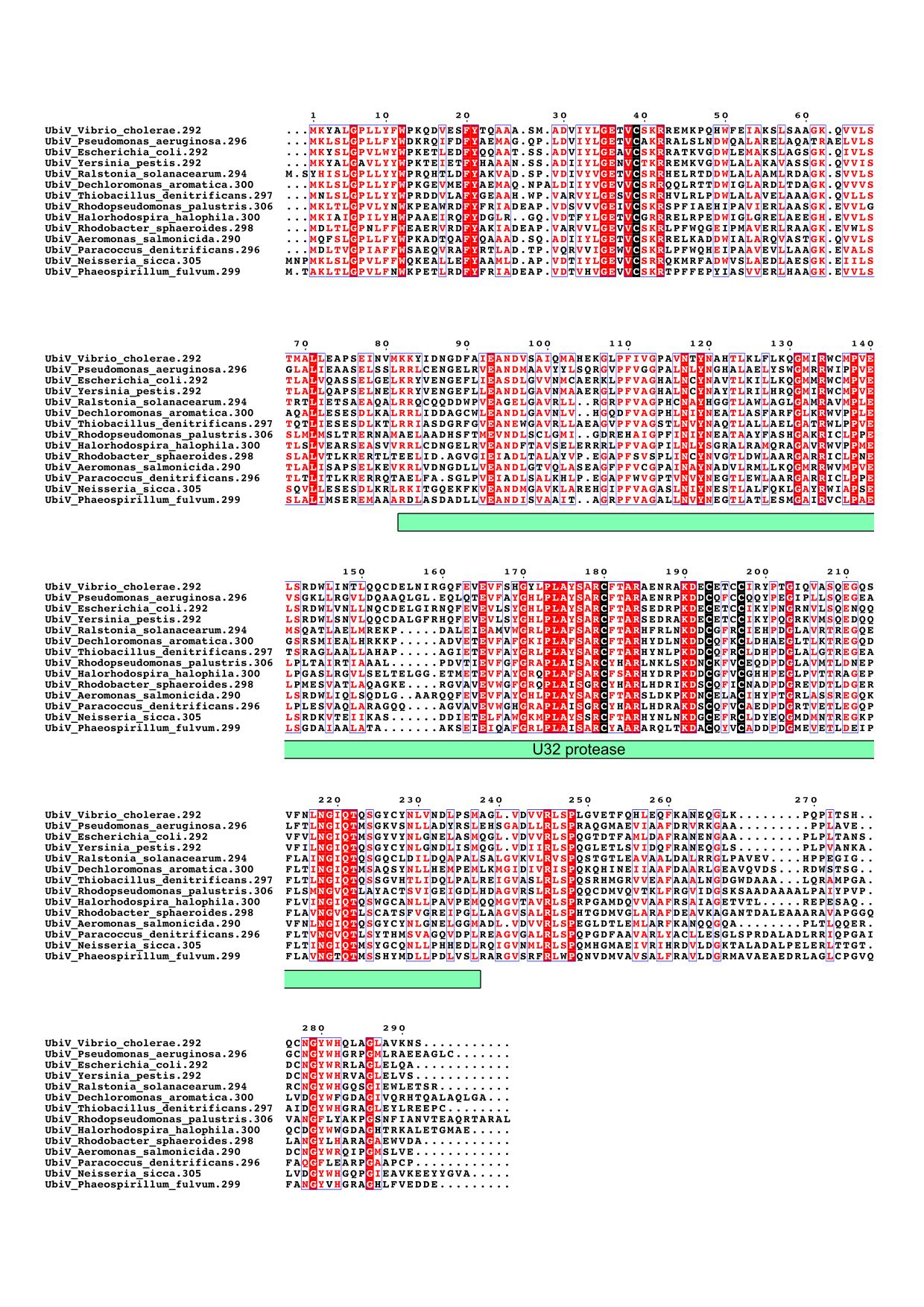

### Supplemental Figure 5

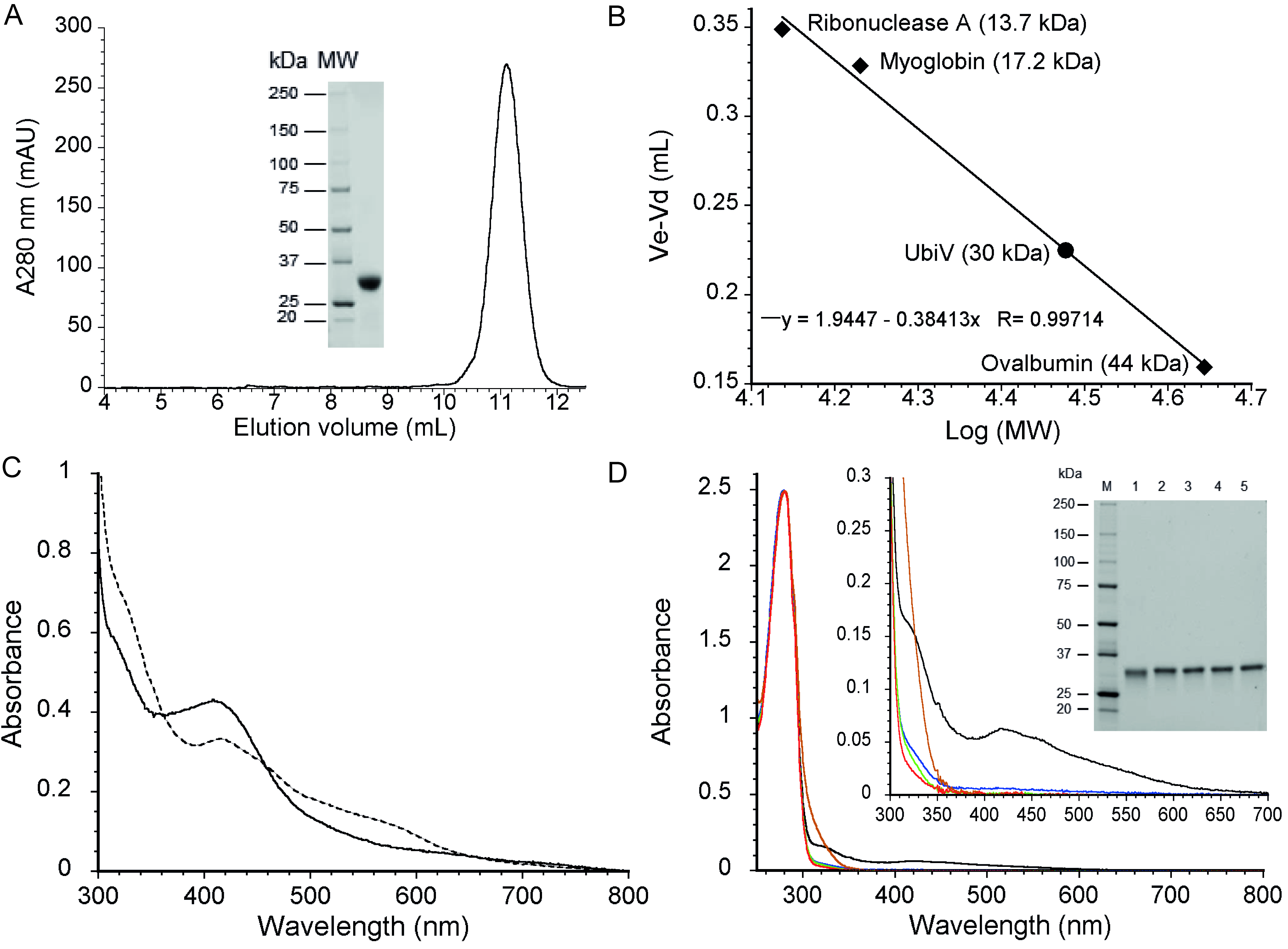

### Supplemental Figure 6

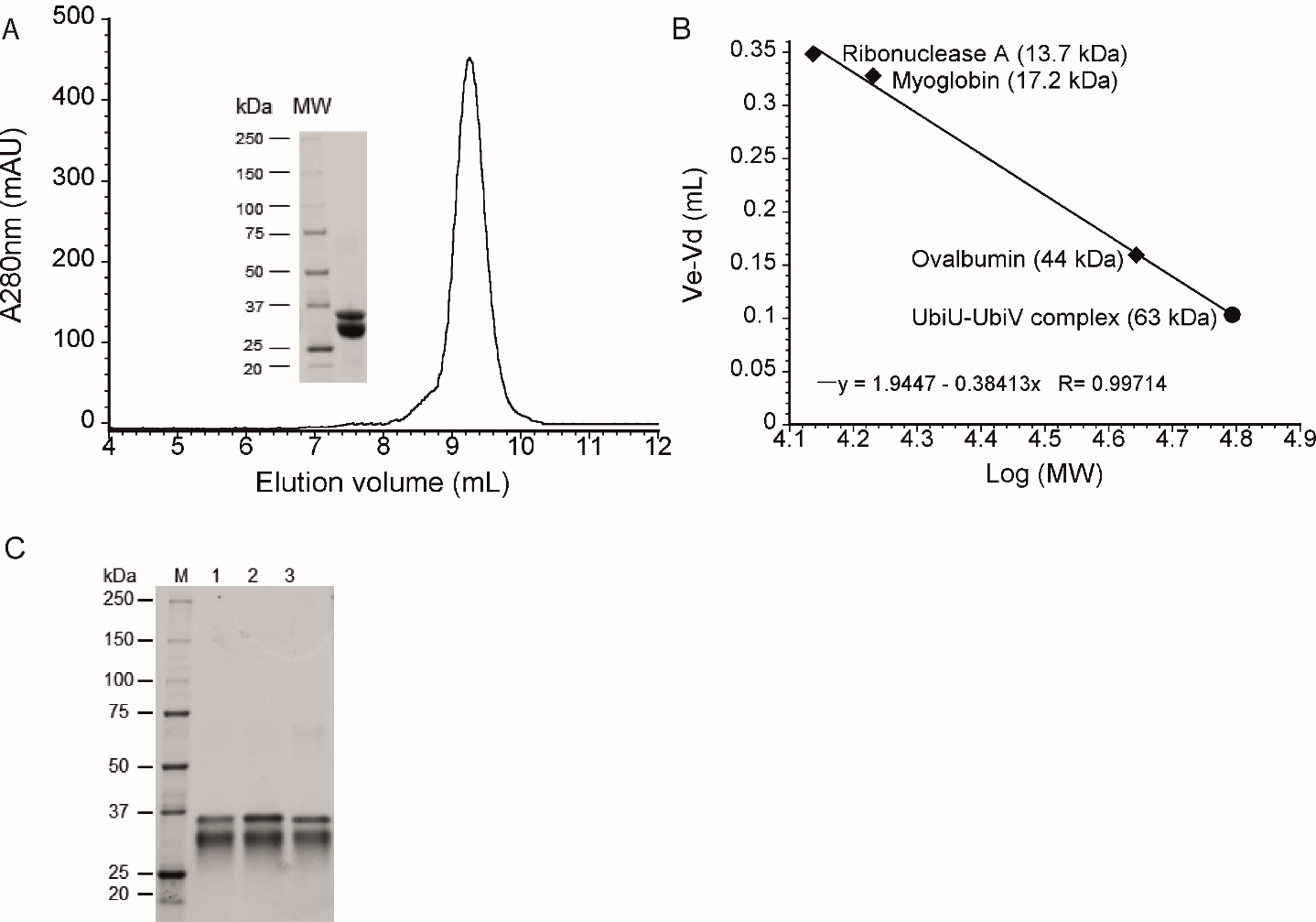
